# BRAF^V600^ and ErbB inhibitors directly activate GCN2 in an off-target manner to limit cancer cell proliferation

**DOI:** 10.1101/2024.12.19.629301

**Authors:** C. Ryland Ill, Nasreen C. Marikar, Vu Nguyen, Varuna Nangia, Alicia M. Darnell, Matthew G. Vander Heiden, Philip Reigan, Sabrina L. Spencer

## Abstract

Targeted kinase inhibitors are well known for their promiscuity and off-target effects. Herein, we define an off-target effect in which several clinical BRAF^V600^ inhibitors, including the widely used dabrafenib and encorafenib, interact directly with GCN2 to activate the Integrated Stress Response and ATF4. Blocking this off-target effect by co-drugging with a GCN2 inhibitor in A375 melanoma cells causes enhancement rather than suppression of cancer cell outgrowth, suggesting that the off-target activation of GCN2 is detrimental to these cells. This result is mirrored in PC9 lung cancer cells treated with erlotinib, an EGFR inhibitor, that shares the same off-target activation of GCN2. Using an *in silico* kinase inhibitor screen, we identified dozens of FDA-approved drugs that appear to share this off-target activation of GCN2 and ATF4. Thus, GCN2 activation may modulate the therapeutic efficacy of some kinase inhibitors, depending on the cancer context.

## Introduction

Developments in therapeutic strategies to treat cancer have led to improved success in the initial stages of cancer treatment^1^, but the acquisition of drug resistance and development of metastasis are roadblocks to improving progression free survival and disease free survival in the clinic^2–6^. With their fast growth rate, rapid adaptive mechanisms, and metastatic phenotypes, lung cancer and melanoma remain two of the most prominent and lethal cancers worldwide^7^.

Melanoma, if not detected and treated early in disease progression, is one of the most deadly cancers due to its fast growth rate and high likelihood of metastasis^8,9^. Over half of all melanomas are driven by a mutation in the BRAF protein at position V600^10^, which leads to constitutive activation of the mitogen activated protein kinase (MAPK) pathway, RAF-MEK-ERK (Fig. S1a). The prevalence of this mutation motivated the development of several targeted kinase inhibitors selective for mutant BRAF over wild-type BRAF^11–13^. Three of these drugs gained FDA approval over the last decade: vemurafenib, dabrafenib, and encorafenib (respective patient C_max_ values of 9.8μM^14^, 1.6μM^15^, and 2.0μM^16^). However, in spite of initial positive and dramatic patient responses to these inhibitors, resistance almost inevitably develops^8,9^.

It is becoming increasingly appreciated that cancer cells use non-genetic adaptive mechanisms to escape the action of cancer drugs^2–6,17,18^. Studies of “persister cells”, cells that survive treatment with targeted cancer inhibitors^3^, and “escapees”, a subpopulation of persister cells that escape quiescence to re-enter the cell cycle in the presence of these drugs^17^, have elucidated that non-genetic adaptive mechanisms are critical to cancer cell survival and eventual acquisition of drug resistance. In melanoma, many of these adaptive mechanisms involve reinitiation of pro-proliferation signaling, alterations in DNA repair pathways, and pro-survival stress signaling^17,19^.

The Integrated Stress Response (ISR, Fig. S1b) is one such pathway that is being highlighted in current non-genetic drug adaptation research^20^. In canonical contexts, the ISR responds to various stresses through activation of four stress-sensing kinases: (1) general control non-derepressible 2 (GCN2) canonically senses amino acid insufficiency via uncharged tRNA^21^; (2) heme-regulated inhibitor (HRI) senses heme deprivation^22^ and mitochondrial stress^23–26^; (3) protein kinase R (PKR) senses cytoplasmic double-stranded RNA^27^; (4) protein kinase R-like ER kinase (PERK) senses un/misfolded protein^28,29^. When activated, each of these dimeric kinases undergoes a conformational change and auto-phosphorylation, and then phosphorylates the translation initiation factor eIF2α, which prevents the binding of GTP to the eIF2α protein^30^. This then lowers the levels of the eIF2-GTP-Met-tRNA complex, which impairs global cap-dependent protein translation^31^, but promotes the translation of mRNAs with unique 5’ untranslated region elements, including ATF4^32^. The ATF4 transcription factor is then translated, highly post-translationally modified, selects a dimerization partner, and translocates to the nucleus where groups of stress-responsive genes are transcribed in a stimulus-specific manner^33,34^. These gene sets can be either pro-survival (including amino acid synthetases, folding chaperones, and DNA repair enzymes)^35–40^ or pro-apoptotic (via induction of CHOP, induction of ATF6, and suppression of Bcl-2 proteins)^39,41^, depending on the strength and duration of the stress^33^.

While GCN2, one of the ISR kinases, is best known for its role in sensing and responding to amino acid limitation^42^, its cellular influence is wider than was initially understood^43^. Recent publications detail GCN2’s role in sensing and regulating stalled or collided ribosomes^44,45^, regulating and being regulated by mTORC1^46–49^, and responding to UV irradiation^50,51^. Further, GCN2 has been implicated in several cancer resistance and metastasis phenotypes^52–55^. Numerous small molecule GCN2 modulators are currently being developed to activate and inhibit GCN2 in hopes of therapeutic benefit in the treatment of cancer, with Hibercell’s HC-7366 recently receiving fast track designation for the treatment of acute myeloid leukemia and renal cell carcinoma^55^.

The ISR is a nuanced and highly variable system, and its activation results in significant transcriptional changes to the cell^56^. Off-target activation of such a system would result in unpredictable, context-dependent outcomes. Over the last three years, four groups have published observations in which the ISR is activated by off-target actions of the kinase inhibitors neratinib, erlotinib, gefitinib, dovitinib, and adavosertib ^57–60^. All describe a mechanism whereby at low doses, the inhibitor binds a single active site of the dimeric GCN2 kinase, which increases affinity of the second active site for ATP, paradoxically activating the kinase^57–59^. As the concentration of the inhibitor increases, both active sites become occluded, and the kinase is then fully inhibited. This results in a characteristic biphasic activation-inhibition curve in response to increasing drug concentration. While the mechanism of off-target GCN2 activation by the above drugs has been described biochemically and structurally, the cellular and tumor-level effects of this off-target are not fully understood.

Herein, we describe the ability of clinical BRAF^V600^ inhibitors dabrafenib, encorafenib, and vemurafenib to elicit this same off-target effect on GCN2. In long-term cancer outgrowth experiments, we show that attenuation of the off-target effect via cotreatment with a GCN2 inhibitor enhances cancer cell outgrowth in A375 melanoma and PC9 lung cancer cell models, suggesting that the off-target effect is detrimental to the cancer cell contexts we tested here.

## Results

### ATF4 induction in response to BRAF inhibitors is uncoupled from MAPK inhibition

Previous work from our lab in melanoma cells identified a subset of drug-tolerant persister cells that initially enter quiescence in response to BRAF/MEK inhibition but escape quiescence within 3-4 days to enter a slow-cycling state (“escapees”)^17,19^. We previously found that escapees express ATF4 and induce an ATF4-dependent transcriptional program^17^. We originally set out to delineate the role and timing of ATF4 induction in escapees under BRAF/MEK inhibition, and thus designed an improved ATF4 activity reporter (UTR.ATF4-mCit, Fig. 1a) by adapting an existing reporter of ATF4 induction^24^ for a fluorescence time-lapse microscopy setting. The sensor is based on a fluorescent protein sequence shielded by the upstream regulatory sequence of ATF4, ensuring translational control mirroring that of the endogenous ATF4 protein (Fig. 1a). By switching out the commonly used RFP for mCitrine, we shortened the half-life of the sensor and reduced protein aggregation, which otherwise problematically activates the PERK-eIF2α-ATF4 axis. We then added a dihydrofolate reductase (DHFR) degron domain to ensure rapid degradation of mCitrine, unless the system is stabilized by addition of trimethoprim (TMP) that masks the DHFR degron, which may be done in a concentration-responsive manner (Fig. S1c-f). We ensured TMP itself did not activate the ISR, using thapsigargin (a known activator of PERK^61^), as a positive control (Fig. S1e). Use of TMP keeps the background ATF4-mCitrine levels low. We then multiplexed our UTR.ATF4-mCit sensor with our DHB-based sensor for CDK2 activity^62^ (to distinguish single-cell proliferation-quiescence decisions, Fig. 1b) and fluorescently labeled H2B^62^ (a nuclear marker for cell tracking) in A375 BRAF^V600E^ human melanoma cells.

**Figure. 1.**
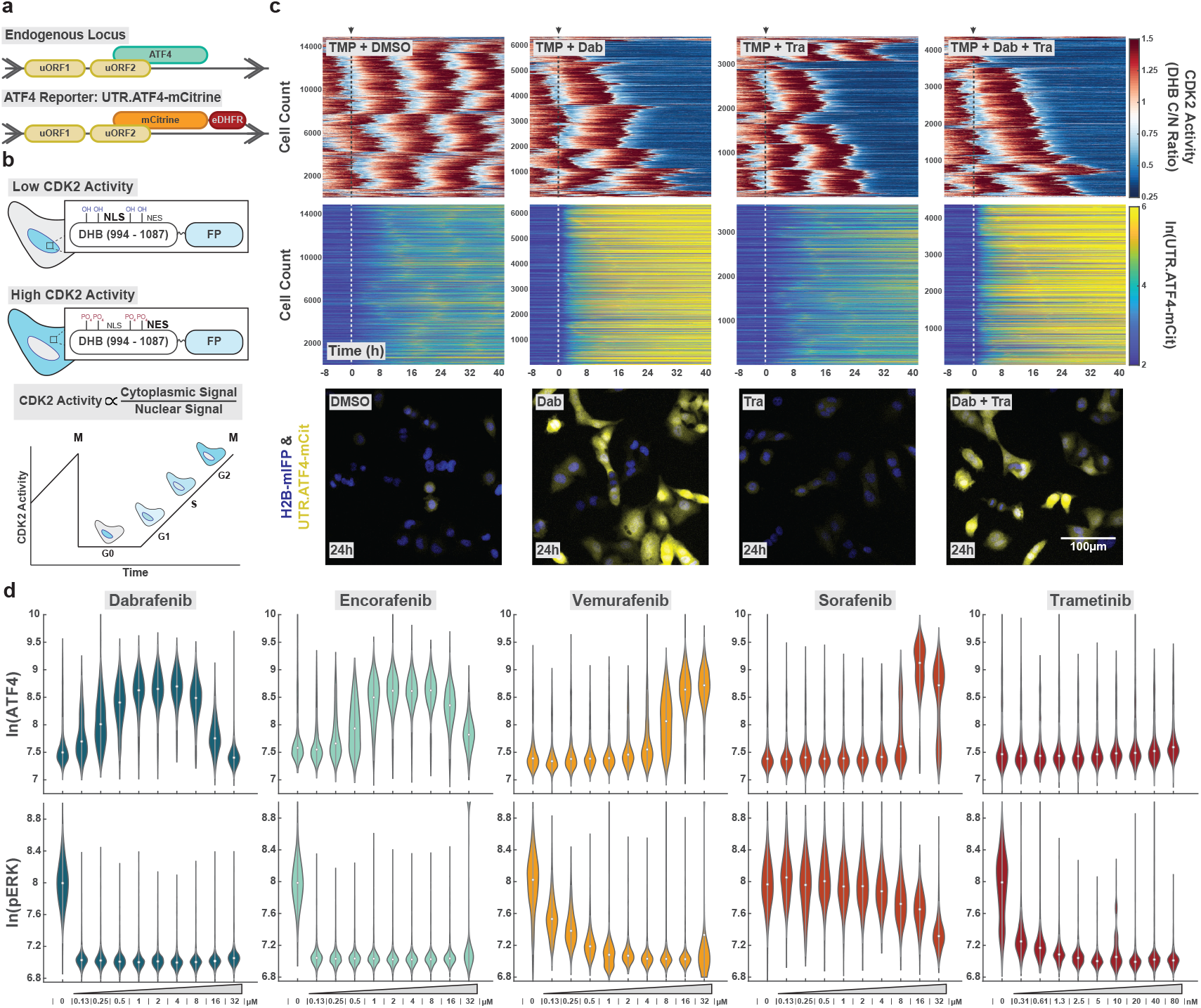
BRAFV600 inhibitors rapidly induce ATF4 in manner that is biphasic with concentration. **a** Schematic of the UTR.ATF4-mCitrine reporter. uORF, upstream open reading frame. **b** Schematic of the DHB CDK2 activity sensor. DHB (DNA-helicase B fragment) localizes to the nucleus when unphosphorylated; progressive phosphorylation leads to translocation of the sensor to the cytoplasm. NLS, nuclear localization signal; NES, nuclear export signal. **c** Three-color A375 H2B-mIFP DHB-mCherry UTR.ATF-mCitrine cells were treated at the dashed line with 1μM trimethoprim (TMP) to stabilize the sensor and 2μM dabrafenib, 10nM trametinib, or the combination and filmed for 48h. Between 3,500 and 14,000 single-cell traces of the CDK2 activity (top) and ATF4 reporter (middle) are plotted in heatmap format according to the colormap. Corresponding representative images of the UTR.ATF4-mCitrine reporter (yellow) and histone 2 B (H2B, blue) are shown below after 24 h of drug. **d** A375 cells were subject to titration of the indicated drug at the given concentrations for 6h, fixed, and stained for ATF4 (top) and phospho-ERK (bottom) and violins of the single-cell nuclear intensities are plotted on a natural log scale (ln).

When we began imaging ATF4 induction under BRAF/MEK inhibition and tracking single cells with EllipTrack^63^, we unexpectedly observed an immediate and intense activation of the UTR.ATF4-mCit sensor in response to dabrafenib treatment in every cell (Fig. 1c column 1-2 and S1c). This immediate ATF4 activation, which began within 1h of treatment, was a distinct phenomenon from what we had previously observed in escapees, which induce ATF4 after 3-4 days of treatment, coincident with cell-cycle re-entry under drug challenge^17^. The rapid ATF4 induction was attenuated by pretreatment with 2BAct (Fig. S1g), an inhibitor that blocks the ISR at the point of eIF2α^64^, indicating canonical ISR signaling via the ISR kinase-eIF2α-ATF4 axis. Immunofluorescence imaging of endogenous ATF4 also revealed a rapid accumulation of ATF4 in response to 2μM dabrafenib with peak ATF4 achieved by 5-6h (Fig S1h). By contrast, inhibition of ERK using 1μM SCH772984 or inhibition of MEK using 10nM trametinib had no effect on ATF4 induction (Fig. 1c column 3 and Fig. S1h), but co-treatment with dabrafenib + trametinib still yielded strong ATF4 induction (Fig. 1c column 4). We conclude that the observed ATF4 induction was not a result of generally inhibiting the MAPK pathway, and hypothesized instead that it was related to dabrafenib itself.

### ATF4 undergoes paradoxical activation-inhibition in response to increasing concentrations of dabrafenib and encorafenib

To determine if dabrafenib exhibited the biphasic activation-inhibition curve previously reported for erlotinib and neratinib^57^, we treated A375 cells with increasing doses of various RAF inhibitors, including dabrafenib (BRAF^V600^), encorafenib (BRAF^V600^), vemurafenib (BRAF^V600^), sorafenib (CRAF), and trametinib (MEK, as a negative control). We found that dabrafenib and encorafenib indeed induced ATF4 in a biphasic manner with peak induction at 2μM and 4μM, respectively (Fig. 1d, top). Importantly, these values are clinically relevant, as the patient C_max_ for dabrafenib is 1.6μM^15^ and the patient C_max_ for encorafenib is 2μM^16^. ATF4 was also induced by high doses of vemurafenib and sorafenib (peak induction 16-32μM), but we did not detect the attenuation of ATF4 at high doses, which would have required testing concentrations beyond the drugs’ aqueous solubility limits. Importantly, trametinib, an allosteric inhibitor of MEK, did not exhibit any induction of ATF4 at any dose tested (Fig. 1d, top). We further found that this biphasic activation-inhibition effect was independent of phospho-ERK level (Fig. 1d, bottom). That is to say, ATF4 could be induced upon blockade of MAPK signaling (dabrafenib, encorafenib, and vemurafenib) or in the presence of MAPK signaling (sorafenib).

### GCN2 activity is necessary for dabrafenib- and encorafenib-induced ATF4 expression

To determine the mechanism by which BRAF^V600^ inhibitors were inducing ATF4, we knocked down each ISR kinase and measured ATF4 induction in response to dabrafenib (Fig. 2a,b). While siRNA methods are commonly used for knockdown, it is well known that double-stranded siRNA/target-mRNA complexes in the cytoplasm can activate PKR^65,66^, the ISR kinase responsible for sensing viral infection. To avoid unintended activation of ATF4, we designed a CRISPRi microscopy platform with single cell resolution and live- and fixed-cell capabilities (scCRISPRi, Fig. S2a,b). We stably expressed the CRISPRi machinery (dCas9–KRAB–MeCP2) with H2B-mTurqoise2 (separated via a P2A sequence) and used flow cytometry to isolate a homogenous population expressing this machinery. Transfection with plasmids containing the sequences for firefly luciferase as well as a sgRNA against our protein of interest allowed us to computationally identify the cells expressing the sgRNA via the luciferase immunofluorescence signal. This strategy allows for classical knockdown comparisons between wells (Fig. 2a) as well as intra-well comparisons, where positive and negative cells from the same well can also be directly compared after computational segregation (Fig. S2c).

**Figure. 2.**
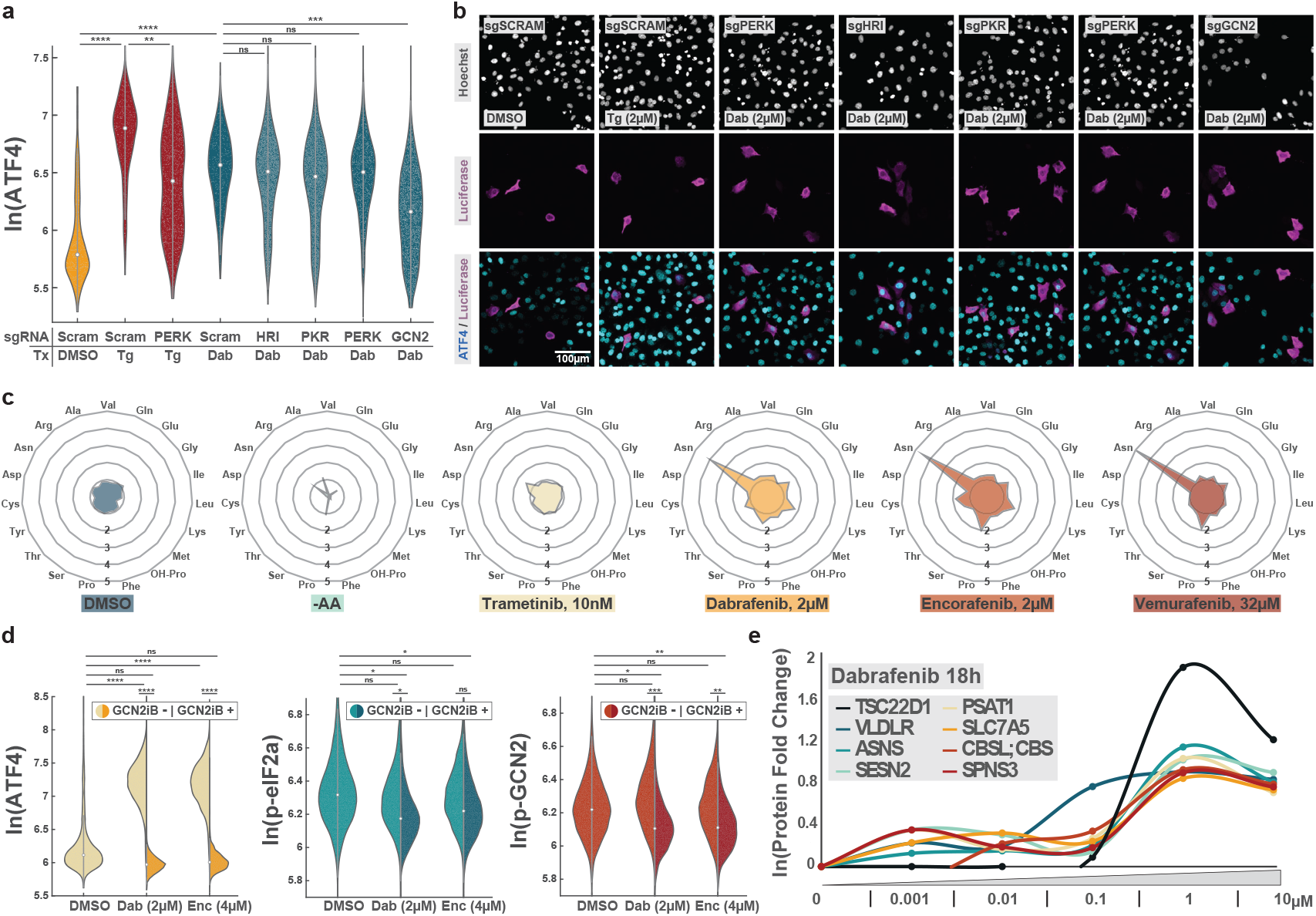
GCN2 is necessary for dabrafenib-induced ATF4 expression. **a** Immunofluorescence of ATF4 in A375 cells plotted as violins after a 48h knockdown of the indicated kinase using scCRISPRi followed by 6h treatment with the indicated compound. The baseline condition (yellow) shows DMSO+transfection with a scrambled guide RNA (Scram). The positive control shows thapsigargin (tg, 1μM) with a scrambled guide RNA (first red violin) or thapsigargin with PERK knockdown (second red violin). The dabrafenib control condition (blue, left, 2μM) is then compared to knockdown of each ISR kinase. Stars represent p-value from two-sample t-test (ns > 10^−7^; * < 10^−7^; ** < 10^−8^; *** < 10^−9^; **** < 10^−10^). **b** Representative images of the quantification shown in (a). Hoechst is shown in grayscale, the luciferase marker protein in magenta, and ATF4 in cyan. **c** Radar plots of fold change of noted amino acid relative to full growth media condition after 6h in the indicated drug. Mean fold change of three independent replicates is shown. Each expanding circle represents an additional fold change from 1 (innermost circle) to 5 (outermost circle). **d** Immunofluorescence quantification of ATF4 (left, nuclear signal), phospho-eIF2a (S51, middle, cytoplasmic signal), and phospho-GCN2 (T899, right, cytoplasmic signal) in A375 cells treated with DMSO, dabrafenib, encorafenib for 6h (left side of split violin), or the given inhibitor in combination with GCN2iB (right side of split violin, 1μM with 1h pretreatment). Stars represent p-value from two-sample t-test (ns > 10^−9^; * < 10^−9^; ** < 10^−10^; *** < 10^−11^; **** < 10^−12^). **e** Top upregulated protein hits by mass spectrometry after 18h of dabrafenib in Jurkat cells at the given concentrations. Lines of best fit of natural log of fold-change were plotted for published fold-change data using the authors’ data browsing application^68^. Of the top 20 upregulated hits, the eight shown displayed biphasic activation-inhibition at ATF4 inducing concentrations.

Employing this scCRISPRi platform, we knocked down each of the ISR kinases for 48h and then treated with 2μM dabrafenib or 1μM thapsigargin for 6h, followed by fixation and immunofluorescence staining (Fig 2a-b). As expected, thapsigargin caused a strong induction of ATF4 that was suppressed by knockdown of PERK. Treatment with dabrafenib also strongly activated ATF4, and this activation was suppressed by knockdown of GCN2 but not by knockdown of the other three kinases. We thus identified GCN2 as the most likely ISR kinase responsible for the observed induction of ATF4.

Since GCN2 is best known for its role in sensing amino acid deprivation^42^, we tested whether dabrafenib was causing a rapid drop in amino acid levels leading to uncharged tRNA and GCN2 activation. We quantified amino acid levels of treated A375 cells using gas chromatography–mass spectrometry, using amino acid-free media as a control. While a 6hr amino acid withdrawal caused a dramatic depletion of most amino acids, as expected, a 6hr treatment with trametinib, dabrafenib, encorafenib, or vemurafenib, did not cause any depletion of cellular amino acids (Fig 2c and S2d-e). Intriguingly, we observe a large 4-5-fold increase in asparagine levels upon dabrafenib, encorafenib, and vemurafenib treatments. Asparagine synthetase (*ASNS*) is one of the best-studied transcriptional targets of the GCN2–eIF2α–ATF4 axis^67^, further supporting that GCN2 is the ISR kinase responsible for ATF4 induction by BRAF^V600^ inhibitors.

To test whether GCN2 kinase activity is required for the induction of ATF4 by BRAF^V600^ inhibitors, we treated A375 cells with 2μM dabrafenib or 4μM encorafenib for 6h together with GCN2iB, an inhibitor of GCN2 (Fig. 2d). Notably, GCN2iB is known to paradoxically activate GCN2 at low concentrations^59^, necessitating use of a high 1μM dose to put GCN2 firmly into an inhibited state (Fig. S2f). Immunofluorescence analysis revealed that dabrafenib or encorafenib co-inhibition with GCN2iB completely blocked induction of ATF4 (Fig. 2d). Co-inhibition with GCN2iB also reduced auto-phosphorylation of GCN2 and phosphorylation of eIF2α, but the results were less strong, likely due to weak antibody signal in this context (Fig. 2d).

In a recent proteomics study, Eckert *et al*. performed titrations of a wide variety of drugs in the Jurkat human T lymphocyte cell line^68^. Intriguingly, of the top 20 most upregulated hits in response to dabrafenib titration, eight of them displayed biphasic activation-inhibition. Among these eight proteins were classic GCN2/ATF4 targets including asparagine synthetase (ASNS), sestrin 2 (SESN2), phosphoserine aminotransferase 1 (PSAT1), and solute carrier family 7, member 5 (SLC7A5)^33^ (Fig. 2e). Additionally, since Jurkat cells do not have the BRAF^V600^ mutation, but rather only wildtype BRAF, dabrafenib is unable to elicit its on-target effects. We therefore hypothesized that GCN2 was being activated in an off-target manner.

### BRAF^V600^ inhibitors activate GCN2 via direct interaction

Since the BRAF^V600^ inhibitors did not spur a canonical activation of GCN2 via amino acid depletion (Fig. 2c), we hypothesized that these drugs may be directly binding GCN2. Three RAF inhibitors had been previously tested against the kinase domain of GCN2 in the LINCS KINOMEscan database (Fig. 3a): dabrafenib, vemurafenib, and sorafenib. Notably, dabrafenib and vemurafenib displayed significant binding affinity for GCN2’s kinase domain at the tested concentration (10μM) (Fig. 3b). Torin1, an inhibitor of mTORC1, is shown as a negative control.

**Figure. 3.**
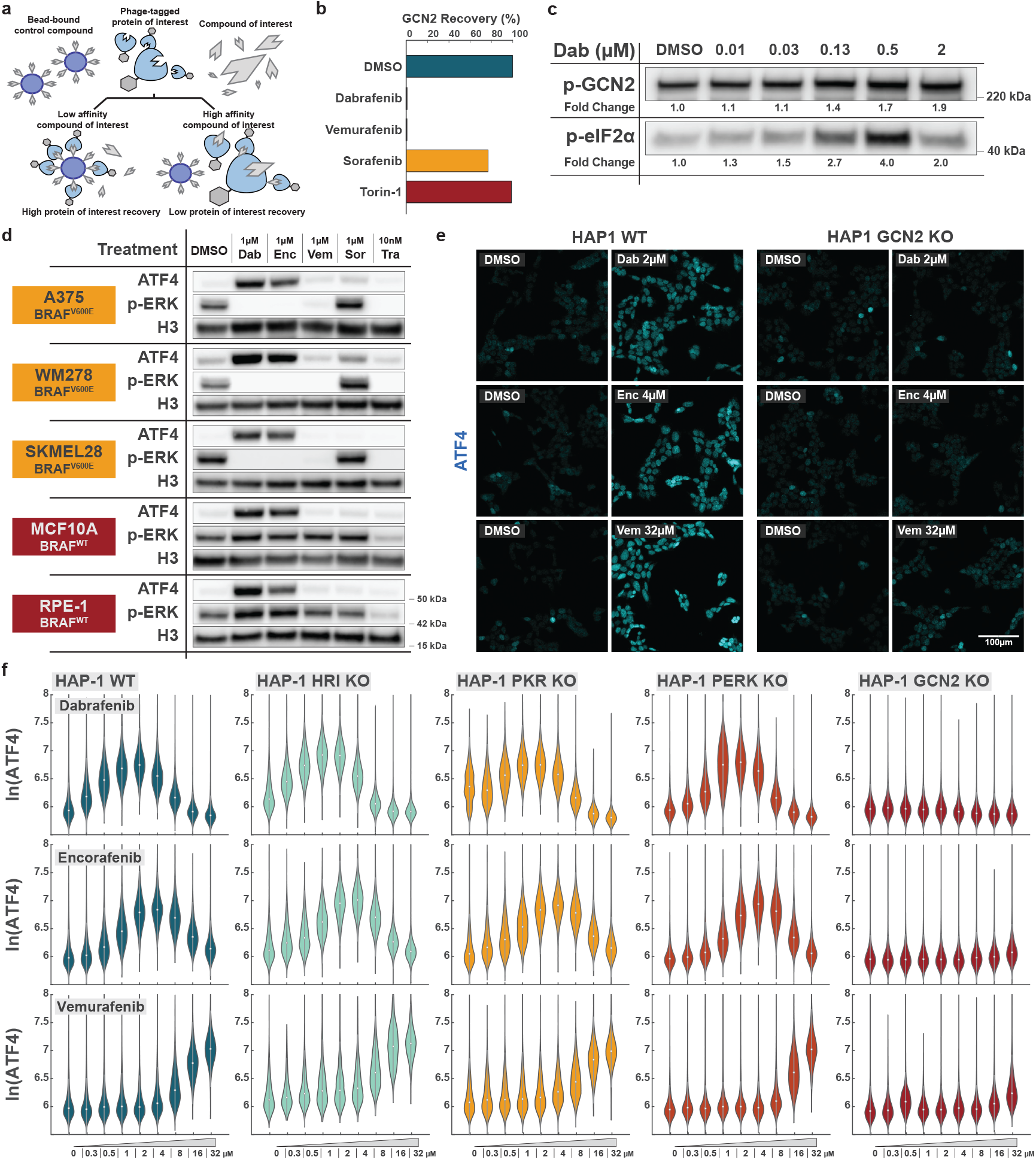
GCN2 is non-specifically activated by RAF inhibitors. **a** Schematic of the LINCS KINOMEscan experimental workflow. This competitive pulldown assay is performed by combining phage-tagged GCN2 kinase domain with a known GCN2 inhibitor bound to agarose beads. A secondary compound is then added. Agarose beads are collected, phage is lysed, and its contents quantitatively sequenced to determine percent recovery. High percent recovery implies low interaction between GCN2 and the secondary compound. Low percent recovery implies high interaction between GCN2 and the secondary compound. **b** LINCS KINOMEscan assay data. Percent GCN2 recovery is plotted. All drugs tested at 10μM. **c** *In vitro* kinase assay of GCN2 and eIF2a phosphorylation in response to increasing concentrations of dabrafenib. **d** The noted drugs were tested in five cell lines: A375 (BRAF^V600E^ melanoma), WM278 (BRAF^V600E^ melanoma), SKMEL28 (BRAF^V600E^ melanoma), MCF10A (non-transformed breast epithelial), and RPE-hTERT (non-transformed retinal pigment epithelial). Cells were treated for 6h, lysed, and probed for the indicated protein species. Histone H3 is used as a nuclear loading control. **e** Immunofluorescence images of ATF4 in WT (left) and GCN2 knockout (KO, right) HAP-1 cells in response to peak activating doses of dabrafenib (2μM), encorafenib (4μM) and vemurafenib (32μM) for 6h. **f** Violins of single-cell ATF4 immunofluorescence intensity in response to titrations of the indicated drug in ISR kinase knockout HAP-1 cell lines as shown in (e).

To further test for a direct interaction with GCN2, we performed an *in vitro* kinase assay with full length GCN2 and eIF2α. Notably, we did not add any tRNA to the reaction mixture. We observed a striking, biphasic phosphorylation of eIF2α, in which lower concentrations of dabrafenib lead to phosphorylation of the substrate, while higher drug concentrations inhibited GCN2’s kinase activity (Fig. 3c). Taken together, these experiments show that a) there is a direct interaction between dabrafenib and GCN2 and b) dabrafenib can bypass GCN2’s native regulatory mechanism to promote its activation without supplemental uncharged tRNA.

We next tested the inhibitors’ abilities to activate ATF4 in a variety of cell lines (Fig. 3d). We included three BRAF^V600E^ melanoma lines (A375, WM278, and SKMEL28) and two non-transformed, BRAF^WT^ lines (MCF10A and RPE-hTERT). Each BRAF inhibitor was tested at 1μM, which is within the ATF4-activating range for dabrafenib and encorafenib, but below the activating threshold for vemurafenib and sorafenib. We reasoned that in the case of a true off-target interaction, we would observe similar ATF4 induction in both the BRAF^WT^ cell lines and the BRAF^V600E^ lines, despite the specificity of dabrafenib, encorafenib, and vemurafenib for the V600 mutant. Indeed, we observed ATF4 induction with both dabrafenib and encorafenib treatment even in BRAF^WT^ cell lines. Additionally, in the BRAF^V600E^ cell lines, 1μM vemurafenib was able to ablate p-ERK signal while only slightly inducing ATF4, again supporting the notion that ATF4 activation is not a direct consequence of MAPK inhibition.

Finally, we measured ATF4 induction by immunofluorescence in HAP-1 chronic myeloid leukemia cells. As a haploid cell line, HAP-1 cells provide a simple genetic background in which to generate CRISPR knockouts of proteins of interest^69^. We treated HRI, PKR, PERK, or GCN2 HAP-1 knockout cells^70^ with increasing concentrations of dabrafenib, encorafenib, and vemurafenib for 6h, and measured endogenous ATF4 levels by immunofluorescence (Fig. 3e,f). All cell lines displayed biphasic activation–inhibition of ATF4 except for the GCN2 knockout cells, which did not induce ATF4 at any concentration of any drug tested (Fig. 3f). We thus conclude that GCN2 is necessary and sufficient for BRAFi-mediated ATF4 induction.

### Structural analysis reveals two distinct binding modes of RAF inhibitors with GCN2

To study the binding modalities of dabrafenib, encorafenib, sorafenib, and vemurafenib to GCN2, we performed computational-based molecular docking studies to facilitate an *in silico* screen. This work revealed two distinct binding modes: an “outer” mode shared by dabrafenib and encorafenib, and an “inner” mode exemplified by sorafenib and vemurafenib (Fig. 4a). These modes were distinguished by differences in active site binding: dabrafenib was predicted to weakly bind Lys^619^ in the GCN2 active site through electrostatic interactions, whereas vemurafenib was predicted to have the same Lys^619^ electrostatic interaction in addition to multiple hydrogen bonds with Cys^805^, Glu^803^, Asp^856^, and Phe^867^ (Fig. 4b). Thus, drugs with the inner binding modality (vemurafenib, sorafenib) would have much higher binding affinities due to increased intermolecular interaction with the binding pocket. Accordingly, sorafenib and vemurafenib were predicted to have significantly higher GCN2 Molecular Mechanics / Generalized Born Surface Area (MM/GBSA)^71^ scores than either dabrafenib or encorafenib (Fig. 4c).

**Figure. 4.**
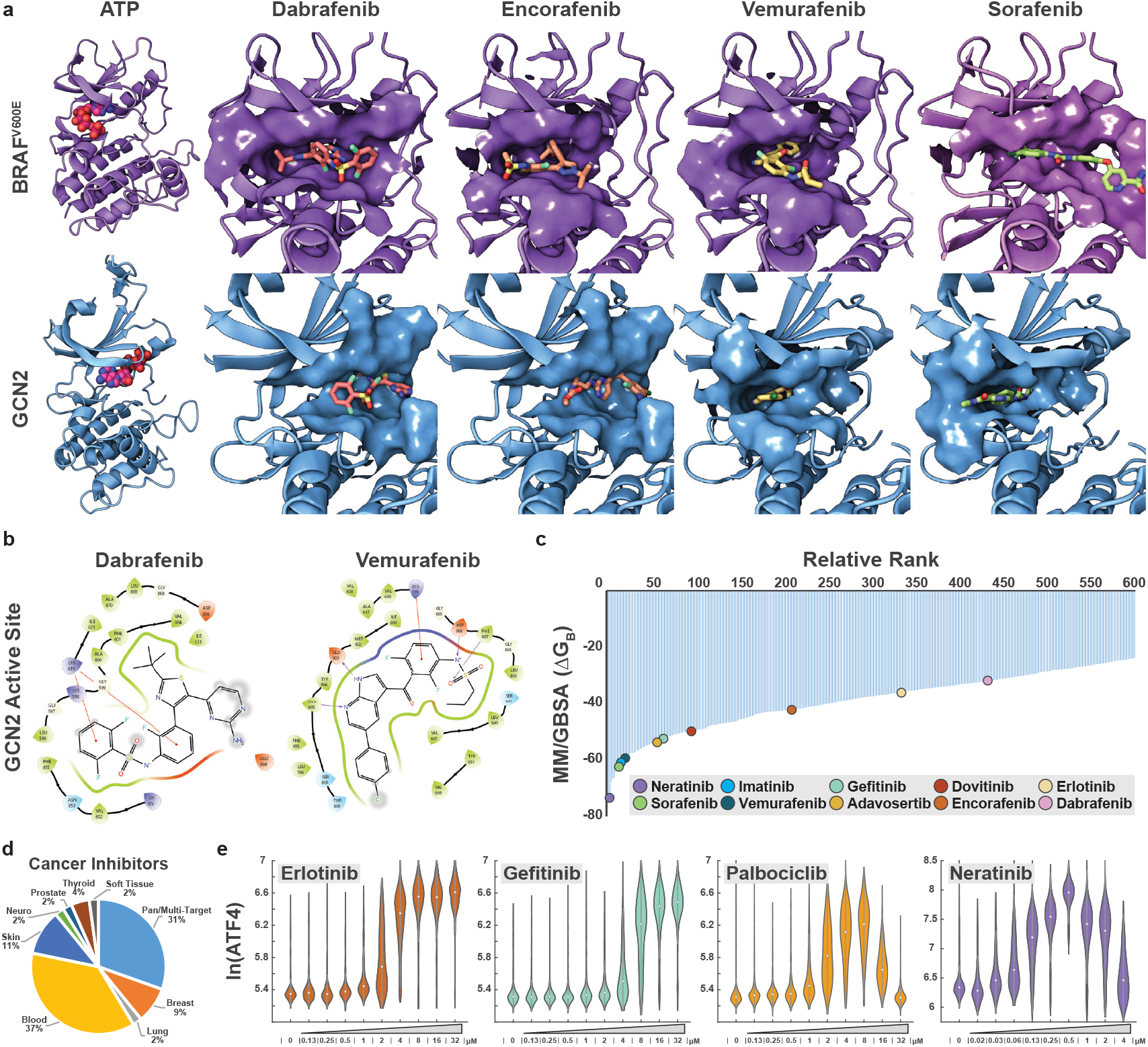
In silico kinase inhibitor screen predicts how RAF inhibitors interact with GCN2. **a** Top: computational docking models of dabrafenib, encorafenib, and vemurafenib bound to BRAF^V600E^ protein, or sorafenib bound to wild-type CRAF. Bottom: the same small molecules’ predicted binding modes to GCN2. **b** Intermolecular interactions of the GCN2 active site with dabrafenib and vemurafenib. **c** Waterfall plot of hits from an in silico kinase inhibitor screen with the kinase domain of GCN2. Hits are ranked by relative MM/GBSA value. **d** Pie chart of the cancer drug targets of the top 200 hits (of which >30% are cancer drugs) in the in silico inhibitor screen. **e** Validation of hits from the in silico inhibitor screen and LINCS database. A375s were treated with the indicated drug for 6h. Immunofluorescence of single-cell nuclear ATF4 signal is shown as violin plots.

While drugs with the inner binding modality may bind with greater affinity, we observe that their activation of GCN2 only occurs at high concentrations. This effect may potentially be due to the kinetics associated with each binding mode – the inner mode is a much stronger interaction, but would require very specific molecular orientations to achieve, leading to a low k_on_, a low k_off_, and high concentration necessary for activation. We would correspondingly expect a tight concentration range over which biphasic activation-inhibition would occur, since a low k_off_ would ensure quick occlusion of both active sites once appropriate concentrations had been reached. That is to say, the stronger the binding mode, the less additional drug needs to be added to bind both of the dimer’s sites and inhibit the kinase due to the low probability of drug dissociation from the initial binding site. In contrast, the outer binding mode (dabrafenib, encorafenib) would not require such specific molecular orientations and may have a correspondingly high k_on_, though the weak interactions of this binding mode would facilitate a high k_off_. In this outer mode, we would expect activation to initiate at lower concentrations but require a much wider concentration range to reach full inhibition, as molecular exchange would be frequent. Indeed, we observe an approximate 120-fold change in concentration necessary for completion of the biphasic activation-inhibition curves of dabrafenib and encorafenib, whereas we can extrapolate an approximate 16-fold delta necessary for completion of the vemurafenib and sorafenib activation-inhibition curves (Fig. 1d).

### *In silico* drug screen suggests that multiple clinical kinase inhibitors may inadvertently activate GCN2

Recent literature has revealed that gefitinib, erlotinib, neratinib, dovitinib, and adavosertib activate GCN2 in the biphasic manner characteristic of the described off-target interaction^57–60^. Coupled with our finding that BRAF^V600^ inhibitors also non-specifically activate GCN2, we assessed the potential breadth of this effect across clinical inhibitors used in the treatment of cancer. By performing an *in silico* kinase inhibitor screen with the Selleckchem FDA-approved & Passed Phase I drug library (3603 compounds), we identified over 400 compounds that are predicted to bind GCN2 with a similar or higher affinity than dabrafenib (Fig. 4c). Sorted by MM/GBSA, vemurafenib and sorafenib both appear in the top 25 hits. In agreement with their differing binding modalities, encorafenib and dabrafenib appear much further down the list (ranks 213 and 434, respectively). Gefitinib, erlotinib, neratinib, and dovitinib, all confirmed activators of GCN2 in cell culture models, are all predicted to bind GCN2 with higher affinity than dabrafenib.

The *in silico* screen reveals hits that include prevalent clinical and research compounds with indications for treating multiple cancers, including compounds utilized to treat and study sarcomas, neuroblastomas, lymphomas, and leukemias (Fig. 4d). We identified and tested the abilities of a number of identified inhibitors to induce ATF4 expression, including erlotinib and gefitinib (EGFR inhibitors used to treat lung cancer), palbociclib (CDK4/6 inhibitor used to treat breast cancer), and neratinib (pan-HER inhibitor used to treat breast cancer) and found they all activate ATF4 in A375 cells (Fig. 4e). Notably, our results in A375 cells indicate that gefitinib and palbociclib do not activate GCN2 at patient-relevant concentrations, while both erlotinib (C_max_ 5.4µM)^72^ and neratinib (C_max_ 152nM)^73^ do. Imatinib and nilotinib also activate GCN2, but not at physiological doses (Fig. S3a). The results of this screen, coupled with published GCN2 off-target data, implicate drugs across the cancer spectrum as potential GCN2 activators.

### GCN2 activation reduces cancer outgrowth and drug resistance

Off-target effects can influence the efficacy of treatment for better or for worse^74,75^. We thus tested the role of the GCN2 off-target effect in cancer cell proliferation. When we examined patient C_max_ values, we noted that dabrafenib and encorafenib are used in the clinic near the peak-activating ATF4 concentrations that we established in cell culture models here, as are erlotinib and neratinib (Fig. 5a). In wondering why so many clinical drugs were used at near-peak GCN2-activating concentrations, we reasoned that higher drug doses would systemically inhibit GCN2 across all of a patient’s cells, which may result in severe generalized toxicity. Therefore, the maximum tolerated dose may, in these cases, occur just before the initiation of GCN2 inhibition.

**Figure. 5.**
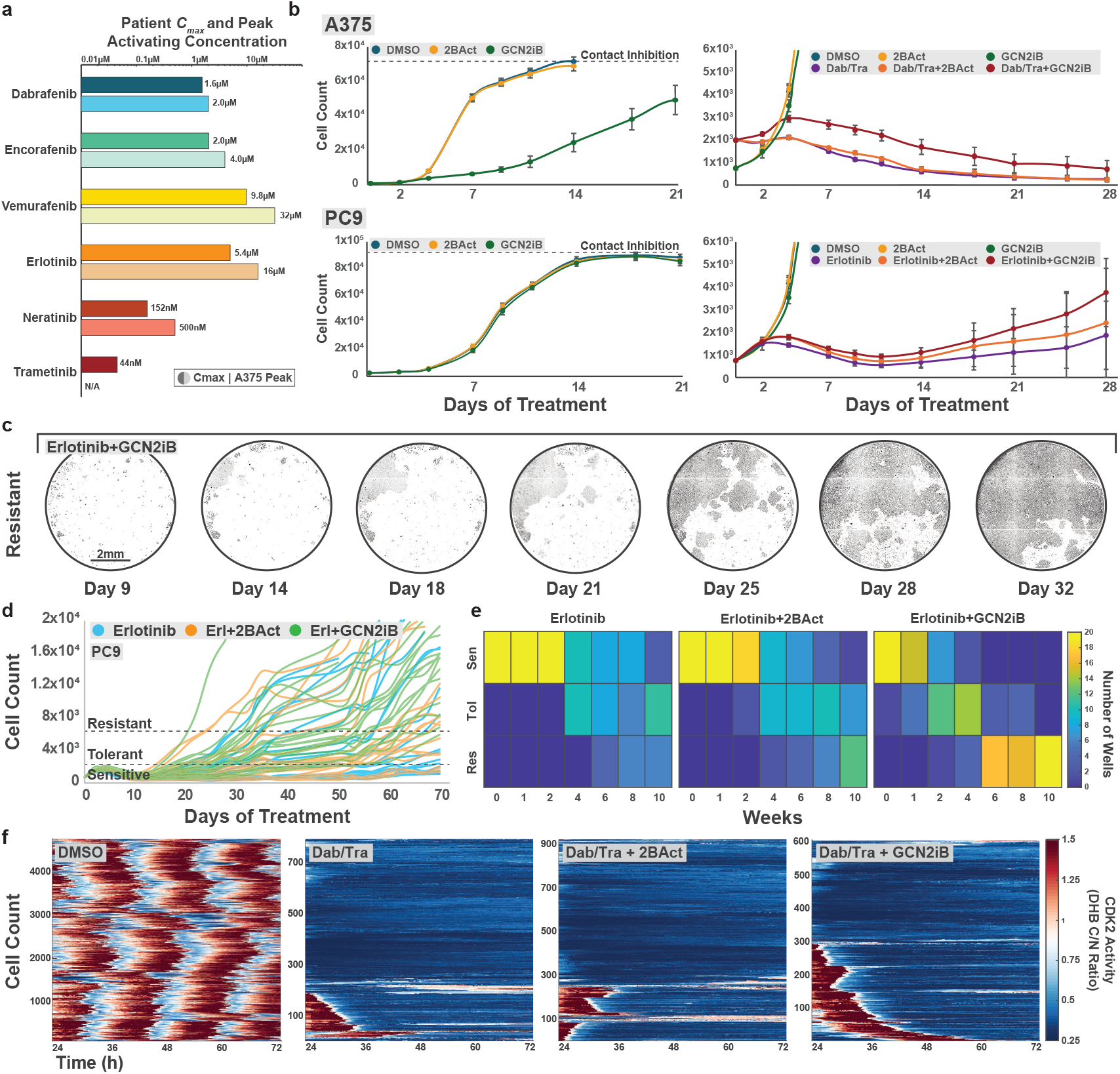
GCN2 inhibition enhances outgrowth and resistance. **a** Mean patient maximum plasma concentrations from the relevant clinical trial for the indicated drug (top bar) compared to peak ATF4-activating concentration (bottom bar). **b** Outgrowth of A375 cells (top) and PC9 cells (bottom) expressing H2B-mCherry was quantified by monitoring cell count via whole-well live fluorescent imaging in 96 well format. Media and drugs were refreshed every 2-3 days. 20 wells per condition were maintained and imaged. Concentrations are as follows: dabrafenib 2μM; trametinib 10nM; erlotinib 5μM; 2BAct 200nM; GCN2iB 1μM. Error bars represent +/-1 standard deviation of replicate wells. **c** Representative whole-well live fluorescent images of PC9 cells treated with erlotinib + GCN2iB that terminate in the “resistant” bin. Further images are available in Fig. S3. **d** Single-well outgrowth traces of PC9 H2B-mCherry cells treated with erlotinib (blue), erlotinib+2BAct (orange), and erlotinib+GCN2iB (green). Cells were arbitrarily binned into sensitive (<1 population doubling), tolerant (greater than or equal to 1 and less than or equal to 4 population doublings), or resistant (>4 population doublings) groups. Traces terminating early were removed from the imaging plate for phenotypic analysis. **e** Heatmaps displaying the number of wells per bin in sensitive, tolerant, or resistant PC9 cells over 10 weeks of treatment in response to the indicated drug condition (erlotinib 5μM; 2BAct 200nM; GCN2iB 1μM). **f** Single-cell CDK2 activity traces of A375 H2B-mIFP DHB-mCherry cells treated with dabrafenib/trametinib, dabrafenib/trametinib + 2BAct, or dabrafenib/trametinib + GCN2iB (dabrafenib 2μM; trametinib 10nM; 2BAct 200nM; GCN2iB 1μM). Cells were treated for 24h prior to imaging, then imaged for 48h.

To test whether the off-target activation of GCN2 was beneficial or detrimental to cancer cell outgrowth in drug, we began long-term treatments of A375^V600E^ melanoma and PC9 EGFR^E746-A750del^ lung adenocarcinoma cells. A375 H2B-mCherry cells were plated in 96-well format and treated with the appropriate kinase inhibitor with or without 2BAct (an inhibitor of ATF4 induction)^64^ or GCN2iB. We performed full-well live fluorescent imaging to monitor cell count over 4-10 weeks non-invasively. In A375 cells, we observed a marked reduction in cell count in response to GCN2iB alone, but not in response to 2BAct alone (Fig. 5b, top left). However, when we combined GCN2iB with the clinical combination of dabrafenib/trametinib, we observed increased cell outgrowth relative to dabrafenib/trametinib alone (Fig. 5b top right and S3b). By contrast, dabrafenib/trametinib paired with 2BAct was not markedly different than dabrafenib/trametinib alone (Fig. 5b, top right).

Perplexingly, we note that non-clinically relevant treatments with dabrafenib+GCN2iB or encorafenib+GCN2iB (without MEK co-inhibition) are significantly more toxic than dabrafenib or encorafenib alone (Fig. S3c-d). We speculate this difference may be due to rapid rebound of MAPK activity (within days) when BRAF^V600^ inhibitors are used as a monotherapy vs. very slow rebound of MAPK activity when BRAF^V600^ inhibitors are combined with MEKi. MAPK reactivation is necessary for mTORC1 reactivation, which we have seen to be important for drug adaptation^17^. The interplay between mTORC1 and GCN2 signaling is complex and not well understood and thus the causes, effects, and extent of the GCN2-mTORC1 interplay remain open questions in the ISR field. These data would suggest, though, that GCN2 signaling can play an anti-proliferative role in a context with minimal mTORC1 activity.

In PC9 EGFR-driven lung cancer cells treated with the EGFR inhibitor erlotinib, we observed the same trend, although these cells were not sensitive to GCN2iB as a monotherapy (Fig. 5b, bottom left). By the 4-week treatment point, wells treated with erlotinib + GCN2iB displayed increased outgrowth and well-to-well variability relative to those treated with erlotinib alone or erlotinib + 2BAct (Fig. 5b, bottom right). We also observed the same trend when testing the pan-HER inhibitor neratinib in PC9 cells (Fig. S3c-d).

We continued monitoring PC9 H2B-mCherry cell count for 10 weeks in each of the treatment conditions (erlotinib, erlotinib+2BAct, erlotinib+GCN2iB). We arbitrarily defined three bins into which we grouped each well at every timepoint: “sensitive wells” (Fig. S3e top) that had not completed a population doubling; “tolerant wells” (Fig. S3e middle) that had completed between one and four population doublings; and “resistant wells” (Fig. 5c and S3e bottom) that had completed over four population doublings (Fig 5d). By plotting the number of wells of each treatment condition in each bin over the duration of the treatment, we observed that wells in the erlotinib + GCN2iB condition maintained significantly more proliferative inertia than those in the other conditions (Fig. 5e). These wells progressed away from the sensitive state (Fig. 5e, top) and towards the resistant state (Fig. 5e, bottom) notably faster than their erlotinib alone or erlotinib+2BAct counterparts. Qualitatively, small islands of resistance can be observed developing and rapidly expanding as resistant cells outcompete those that maintained sensitivity within the same well (Fig. 5c and S3e).

We performed live-cell timelapse imaging of CDK2 activity to determine the acute cell-cycle effects of GCN2 inhibition on A375 cells treated with dabrafenib/trametinib (Fig. 5f). We observed a higher fraction of cycling cells under dabrafenib/trametinib+GCN2iB cotreatment conditions compared to cells treated with dabrafenib/trametinib alone or cotreated with 2BAct, mirroring the data in Fig. 5e. We found that GCN2-inhibited cells were more likely to remain in the cell-cycle after the first day of dabrafenib/trametinib treatment, and more likely to re-enter the cell cycle on the second day of dabrafenib/trametinib treatment than cells treated with dabrafenib/trametinib alone. These data suggest that the outgrowth observed in GCN2iB-cotreated cells is driven by a higher proliferative rate. GCN2 co-inhibition may thus enable the development of resistant phenotypes by promoting cell cycling and passage through S phase where many mutations occur.

## Discussion

Herein, we describe the biochemical and biological impacts of an off-target effect of multiple cancer inhibitors on the dimeric GCN2 kinase. Using scCRISPRi, genetic knockout cell lines, GC-MS, *in vitro* kinase assays, and analysis of published data sets, we show that GCN2 is responsible for the observed ATF4 activation via a direct, off-target interaction between the BRAF^V600^ inhibitors and the kinase. Our results agree with the published mechanism in which at low concentrations, the drug competitively interacts with a single active site of the GCN2 dimer. This interaction induces a conformational change which results in an increased affinity for ATP in the dimer’s second active site, triggering the kinase’s catalytic activity. As drug concentration increases, both active sites become occluded, leading to inhibition of the dimer^57–59^. In response to increasing concentrations of drug, this mechanism results in a characteristic biphasic activation-inhibition curve of GCN2 activity, which we measure by ATF4 expression. Direct GCN2 *in vitro* kinase assays mirrored the results of our indirect ATF4 induction assays by displaying the same biphasic pattern in phosphorylation of eIF2α, GCN2’s best-characterized substrate that is directly responsible for ATF4’s translational regulation. We further examined the binding of these inhibitors with GCN2’s kinase domain computationally, during which we discovered two distinct binding modalities of these inhibitors to GCN2. Importantly, it appears that the binding modality employed by a particular drug may be predicted from its GCN2-activating concentration. We observed that drugs predicted to use the outer binding mode (dabrafenib, encorafenib) activated GCN2 at lower peak concentrations, but over a much broader concentration range, most probably due to weaker intermolecular forces between the drug and binding site. Alternatively, drugs employing the inner binding mode (vemurafenib, sorafenib) activated GCN2 at much higher peak concentrations, but likely over a much narrower concentration range. Correspondingly, these binding events were predicted to have much stronger intermolecular interactions. Notably, an *in silico* inhibitor screen revealed dozens of FDA-approved compounds that appear to share this off-target effect on GCN2 at clinically relevant doses. Of the screen hits that we attempted to validate, all showed some degree of GCN2-activating behaviors.

In the last few months, two collaborative bioRxiv preprints also identified an interaction between RAF inhibitors and GCN2^76,77^. Via mutational and structural studies, these publications detail the chemical interactions between RAF inhibitors and GCN2, as well as the conformational and affinity changes to GCN2 upon drug binding. Taken together, our combined work to this point establishes and confirms this potent off-target effect, and provides the cell-based results needed for future *in vivo* studies of the off-target effect on cancer growth and progression.

Here, we tested the consequences of the off-target activation of GCN2 on A375 melanoma and PC9 lung adenocarcinoma cell culture models in terms of cell outgrowth. We found that co-inhibiting GCN2 (and thus mimicking the elimination of the off-target effect) caused increased proliferation in A375 and PC9 cells, and cell outgrowth and resistance in PC9 cells. This finding is in agreement with recent publications on GCN2-activating drugs^55^ and off-target activation of GCN2 by the Wee1 inhibitor adavosertib^60^. In that work, *in vivo* CRISPR screening revealed that off-target GCN2 activity sensitizes cancer cells to the anti-proliferative effects of the Wee1 inhibitor^60^. Similarly, GCN2-activating agents are showing promise in clinical trials to limit tumor growth and promote tumor clearance^55^.

According to the literature surrounding this off-target effect and our *in silico* drug screen results, off-target activation of GCN2 may be unexpectedly widespread across multiple cancer drugs used in multiple cancer contexts. As one of our most ancient and evolutionarily-conserved kinases^78^, we speculate that GCN2 may lack the levels of specificity afforded to more modern kinases, allowing for more promiscuous binding of drugs with varying chemical structures. The breadth of this off-target GCN2 activation across cancer drugs presents interesting therapeutic opportunities. While there may be contexts in which GCN2 activation in cancer cells is detrimental to the patient, GCN2 activity presents as beneficial in the drug and cancer contexts tested here. Activating GCN2, then, may be therapeutically beneficial in such cancer contexts.

Modulating GCN2 activity, though, is not a trivial endeavor. Because of the biphasic activation-inhibition activity observed with GCN2iB, the contrasting cellular outcomes when GCN2 is activated or inhibited, and the fact that dabrafenib and encorafenib reach most normal cells in the body in addition to melanoma cells, this off-target effect may be limiting the patient maximum tolerated drug dose. Indeed, the doses used in the clinic for dabrafenib, encorafenib, and erlotinib suggest that patient benefit is achieved at doses *lower* than the GCN2-inhibitory dose, but within the activating region. In the clinic, dabrafenib and encorafenib are combined with MEK inhibitors (trametinib and binimetinib) to more fully block the MAPK pathway and improve efficacy^79^. Patients benefit from increased MAPK inhibition, but not at the cost of GCN2 inhibition. Dabrafenib or encorafenib, therefore, cannot be dosed higher, and a secondary, non-GCN2-activating inhibitor is added to the treatment (trametinib or binimetinib). It may therefore be a beneficial therapeutic approach to combine targeted kinase inhibitors that *do not* bind GCN2 with specific GCN2 activators (such as HC-7366, currently in clinical trials). With this strategy, the targeted kinase inhibitors could potentially be dosed higher, increasing on-target impact, while maintaining a GCN2-activation phenotype.

In recent months, a pan-RAF inhibitor, tovorafenib, has shown promising phase I results in BRAF-mutant melanoma^80^. Notably, tovorafenib does not need to be combined with a MEK inhibitor, such as trametinib, to achieve these outcomes. We have shown that tovorafenib does not induce ATF4 at patient-relevant concentrations (C_max_ 9μM) in A375 cells (Fig. S3f). We speculate that the higher dosing of this RAF inhibitor (perhaps partly due to lack of a GCN2 off-target effect) relative to clinical standards dabrafenib and encorafenib may be contributing to its clinical effects. Based on our results, it would be worth testing whether tovorafenib in combination with a GCN2 activator provides an even more robust therapeutic response. If this concept shows promise, we would posit that drug design should test for and exclude GCN2 binding to allow space for a separate GCN2 activator in a combination treatment approach. Based on findings from our computational screen, this approach could potentially benefit multiple cancer types treated with targeted kinase inhibitors.

While prevalent in this paper as a marker of GCN2 activation, we note that ATF4 itself does not appear to be greatly affecting the outgrowth and resistance observed based on the fact that inhibiting ATF4 expression using 2BAct (Fig. S1f) did not affect A375 outgrowth with dabrafenib/trametinib and only slightly affected outgrowth in PC9 cells with erlotinib (Fig. 5b,d-f). This finding is in line with a current shift in thinking in the ISR field. While GCN2 is canonically known as the amino acid deprivation sensor of the ISR, the field is beginning to understand an additional role as a direct modulator of mTORC1 activity independent of ATF4 signaling^46–49^. We and many others have demonstrated the importance of mTORC1 signaling in cancer proliferation, drug adaptation, resistance development, and metastasis^17,81,82^. With the direct interplay of GCN2 and mTORC1 signaling coming to light, as well as the identification of the GCN2 off-target effect in several drugs across the cancer drug spectrum, we speculate that GCN2 may influence cancer outgrowth and resistance in an mTORC1-dependent manner. The mechanisms of such an effect remain unclear, though, and detailed signaling studies remain necessary. Future efforts will also be needed to study the *in vivo* effects on cancer progression of enhancing or suppressing the off-target activation of GCN2.

## Acknowledgements

We would like to thank the members of the Spencer Laboratory for general support, help, and discussion, Roy Parker for providing us with HAP-1 ISR kinase knockout cell lines, Theresa Nahreini and the cell culture facility for cell sorting (RRID:SCR_018988), and Joe Dragavon of the BioFrontier Institute’s Advanced Light Microscopy Core (RRID: SCR_018302). This work was performed with the experience, aid, and resources of the BioFrontiers Computing Core at the University of Colorado BioFrontiers Institute. The Revvity Opera Phenix is supported by NIH grant 1S10OD025072. The Aria Fusion FACS sorter is supported by NIH grant S10OD021601. AMD was supported by a Jane Coffin Childs fellowship and a fellowship from the Charles A King Trust. MGVH is supported by the Ludwig Center at MIT, the MIT Center for Precision Cancer Medicine, and the NCI (R35CA242379, P30CA014051). This work was supported by awards from the Damon Runyon Cancer Research Foundation (DRR-68-21) and The Mark Foundation for Cancer Research (DRR-68-21) to S.L.S.

## Author Contributions

C.R.I. performed the majority of the experimental design, experiments, analyses, data interpretation, data presentation, and drafted the initial manuscript. N.C.M created and validated the UTR-ATF4-mCit sensor, performed time-lapse imaging, titrations and corresponding immunofluorescence, and outgrowth imaging and analysis. V.Nguyen and P.R. provided the *in silico* docking and screening studies. V.Nangia performed time-lapse analysis and contributed to experimental design. A.M.D. and M.G.V.H provided the gas-chromatography mass-spectrometry experiment and analysis. S.L.S. contributed to project design, experimental design, data interpretation, and wrote the manuscript with C.R.I.

## Methods Cell culture

A375 BRAF^V600E^ human melanoma cells (#CRL-1619) were purchased from American Type Culture Collection (ATCC). A375 cells were cultured in DMEM (Thermo Fisher, #12800-082) supplemented with 10% FBS, 1.5 g/L sodium bicarbonate (Fisher Chemical, #S233-500), and 1X penicillin/streptomycin, and grown at 37°C with 5% CO_2_. The WM278 human melanoma cell line was obtained from Dr. Natalie Ahn (University of Colorado Boulder). The SKMEL28 human melanoma cell line was obtained from Dr. Neal Rosen (Memorial Sloan Kettering Cancer Center). WM278 and SKMEL28 cells were cultured in RPMI1640 (Thermo Fisher, #22400-089) supplemented with 10% FBS, 1X Glutamax, 1X sodium pyruvate (Thermo Fisher, #11360-070) and 1X penicillin/streptomycin. MCF10A human breast epithelial cells (#CRL-10317) were purchased from ATCC. MCF10A cells were cultured in DMEM/F12 supplemented with 5% horse serum, 20ng/ml EGF, 10ug/ml insulin, 0.5 ug/ml hydrocortisone, 100ng/ml cholera toxin, and 1X penicillin/streptomycin. RPE-hTERT (#CRL-4000) were obtained from ATCC and grown in DMEM/F12 supplemented with 10% FBS, 1x Glutamax, and 1X penicillin/streptomycin. HAP-1 human chronic myeloid leukemia cell lines (wild-type and knockout) were obtained from and validated by Dr. Roy Parker (University of Colorado Boulder)^70^. HAP-1 cells were cultured in the same manner as A375 cells.

### Cell line generation

A375 cells were transduced with H2B-mIFP, DHB-mCherry, and UTR.ATF4-mCitrine lentivirus (Fig. 1 and S1), with CRISPRi-P2A-H2B-mTurqoise2 (Fig. 2 and S2), with H2B-mCherry lentivirus (Fig. 5b-d and S3), or with H2B-mIFP and DHB-mCherry lentivirus (Fig. 5f) as described previously^62^. Cells stably expressing these sensors were isolated by fluorescence activated cell sorting (FACS). Single cell clones of ATF4-sensor cells and CRISPRi cells were acquired by limiting dilution after a bulk FACS sort.

### Small molecules

Small molecules used in this study were: dabrafenib (MedChemExpress, #HY-14660), encorafenib (Selleckchem, #S7108), vemurafenib (Selleckchem, #S1267), sorafenib (Selleckchem, #S7397), trametinib (MedChemExpress, #HY-10999), TMP (ThermoFisher #AAJ6305303), thapsigargin (Sigma-Aldrich, #0000192053), GCN2iB (MedChemExpress, #HY-112654), erlotinib (MedChemExpress, #HY-50896), gefitinib (Selleckchem, #S1025), palbociclib (Selleckchem, #S1116), neratinib (MedChemExpress, #HY-32721), 2BAct (Cayman Chemicals, #37788), imatinib (MedChemExpress, #HY-15463), nilotinib (MedChemExpress, #HY-10159), tovorafenib (MedChemExpress, #HY-15246).

### Antibodies

Primary antibodies and dilutions used in this study include: ATF4 clone D4B8 (Cell Signaling Technologies, #11815, 1:500 immunofluorescence, 1:1000 western blot), phospho-ERK clone D13.14.4E (ERK1/2, Thr202/Tyr204, Cell Signaling Technologies, #4370, 1:1000 immunofluorescence, 1:1000 western blot), phospho-eIF2α clone D9G8 (Ser51, Cell Signaling Technology, #3398, 1:1000 immunofluorescence), phospho-GCN2 clone E1V9M (Thr899, Cell Signaling Technology, #94668S, 1:1000 immunofluorescence), histone H3 clone 96C10 (Cell Signaling Technology, #3638S, 1:2000 western blot). Secondary antibodies and dilutions used in this study include: goat anti-rabbit or goat anti-mouse immunoglobulin G (IgG) secondaries linked to Alexa Fluor 546 (Thermo Fisher Scientific, #A-11010 and #A-11003, 1:1000 immunofluorescence), Alexa Fluor 647 (Thermo Fisher Scientific, no. A-21245 and no. A-21235, 1:1000 immunofluorescence), or horseradish peroxidase (HRP) (Cell Signaling Technology, #7074 and #7076, 1:5000 western blot).

### Immunofluorescence

Cells were seeded on glass-bottom 96-well plates (Cellvis, #P96-1.5H-N) coated with collagen (Fisher Scientific, #5005-100mL, 1:50 in PBS for 20-30m at 37°C). −48h before drug treatment. For experiments involving co-treatment with GCN2iB or 2BAct, cells were pre-treated with those drugs for 1h prior to addition of the primary drugs. All titration experiments were performed with a treatment time of 6h. Plates were fixed/permeabilized with methanol for 20m at −20°C. Blocking was performed for 1h at room temperature in 3% BSA/PBS (w/v) solution. Primary antibodies were incubated overnight at 4°C in 3% BSA/PBS solution. Secondary antibodies were incubated for 2h at room temperature in 3% BSA/PBS solution. Nuclear staining was performed with Hoechst 33342 for 10m in PBS (1:10,000 in PBS; Thermo Scientific #H3570). All immunofluorescence images were obtained on a Nikon Ti-E microscope equipped with a 10X 0.45 numerical aperture (NA) objective. Immunofluorescence signals were quantified as previously described^19^ and as follows: ATF4 (mean nuclear signal), p-eIF2α (mean cytoplasmic signal), p-GCN2 (mean cytoplasmic signal).

### Single-cell CRISPRi

scCRISPRi was performed using A375 cells expressing catalytically inactive Cas9 (dCas9) fused to KRAB and MeCP2 inhibitory domains driven off of the CMV promoter. H2B fused to mTurqoise2 was driven off of the same promoter and separated via P2A peptide. Cells were sorted by FACS into bulk positive populations, and then plated as single cells by limiting dilution. Individual wells were expanded into clonal populations that were used in experiments.

scCRISPRi cells were then transfected with plasmids containing sgRNAs (see below, all sequences sourced from Guo *et al*.^24^) targeting the genetic sequence of the protein of interest driven off of the U6 promoter. The same plasmids contained a second hPGK promoter from which firefly luciferase was expressed. Transfection was performed using Fugene 6 (Promega #E2693). Per 10?l reaction, we used: 1.2?l of Fugene 6, 0.2?g of relevant plasmid, and Opti-MEM media (Fisher Scientific #31-985-070) to 10?l. This 10?l reaction was added to 90?l of full growth media (FGM, see *Cell Culture*) in one well of an imaging plate containing 2000-3000 pre-seeded A375 scCRISPRi cells. Non-targeting, or “scramble,” sgRNAs were used as transfection controls. Cells were incubated in this reaction mixture at 37°C and 5% CO_2_ for 48h, then treated with the relevant drug course. By transfecting in FGM, we maintained cell viability while ensuring low transfection efficiency, allowing for computational comparisons of transfected and non-transfected cells. After treatment, the above immunofluorescence protocol was employed. ATF4 and luciferase stains were multiplexed, allowing for identification of ATF4 levels in knockdown-positive and knockdown-negative cells.

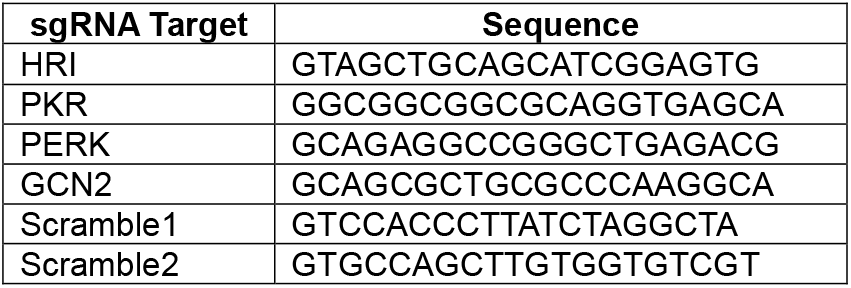

### Live-cell time-lapse imaging and processing

Cells were seeded on glass-bottom 96-well plates coated with collagen 24-48h before drug treatment as described above. Movie images were taken on a Nikon Ti-E using a 10X 0.45 NA objective with filter sets appropriate to the employed fluorescent proteins. Images were obtained every 15 minutes. Imaging was performed in a humidified incubation chamber at 37°C with 5% CO_2_. Phenol red-free FGM (Corning, #90-013-PB) was employed for time-lapse imaging (“imaging media”) to limit background fluorescence. Time-lapse images were computationally processed using our published EllipTrack software package, as previously described^19,63^.

### Live-cell outgrowth imaging

Cells were seeded on plastic-bottom 96-well plates (Revvity #6055500) in imaging media. Drug treatment was started on Day 0, and fluorescent imaging using the Revvity Opera Phenix (5X 0.16 NA) was performed on Days 2, 4, and 7, then every 3 or 4 days thereafter. Media changes/drug refreshes were performed every 2 or 3 days (always immediately before imaging to remove debris). Cell count was determined using Revvity’s onboard nuclear masking software, in which nuclei were masked based on H2B-mCherry signal, and the total number of nuclear masks was quantified.

### Statistics

For the violin plots, the median of the data is indicated by a white circle. The 25^th^ through 75^th^ percentile range is indicated with a light gray bar through the center of the violin. The full data distribution is indicated by the complete shape of the violin, with width corresponding to data frequency. Data plotted throughout are representative of at least two independent experiments with at least two technical replicates per experiment (see supplementary data document detailing the number of replicates for each experiment). Statistical testing was conducted using MATLAB (violin analysis, two-sample t-tests) and Excel (standard deviation error bars).

### Western blotting and *in vitro* kinase assay

Cells were harvested and incubated in ice cold RIPA lysis buffer with 1X phosphatase and protease inhibitors (MilliporeSigma, #4906845001 and #5892970001) for 20m. Lysates were resuspended by vortex, and subjected to sonication for 2m. Samples were centrifuged at 20,000rcf at 4°C for 20m. Supernatant was harvested. Protein content was determined via Pierce BCA Protein Assay Kit (#23227). Lysates were prepared for SDS-PAGE analysis at 20µg per sample using NuPAGE LDS Sample Buffer (#NP0007). SDS-PAGE was run using Bolt 4-12% Bis-Tris Plus gels (Thermo Fisher Scientific, #NW04125BOX). Western blots were performed by transferring to iBlot 2 PVDF Regular Stacks (Thermo Fisher Scientific, #IB24001) using the iBlot 2 transfer system. Membranes were blocked overnight in 0.1X PBS/Casein (Bio-Rad, #1610783). Primaries were incubated overnight at 4°C in 0.1X PBS/Casein with 0.1% Tween-20 (Fisher, #BP-337100). Secondaries were incubated for 2h at room temperature in 0.1X PBS/Casein with 0.1% Tween-20. Blots were developed with Radiance Plus (Azure, #AC2103) substrate and chemiluminescent signal was imaged on an Azure C600.

GCN2 protein and eIF2α protein were purchased from Millipore Sigma (#14-934 and #SRP5232-50UG). One complete *in vitro* kinase reaction is as follows: 20nM TRIS pH 7.5, 200nM EDTA, 0.1% Tween-20, 0.5% glycerol, 10mM MgCl_2_, 50µM ATP, 0.1µg GCN2 kinase domain, 1µg eIF2α, indicated drug concentration, and water to 30µl total volume. The reaction was run for 30min at 37°C. A quench was performed with 10µl of 200mM EDTA. 13.3µl of 4X LDS + DTT (200mM for 50mM final) was then added, and the reaction boiled at 95°C for 10m. The above western blot protocol was then followed.

### Gas-chromatography mass-spectrometry

A375 cells were cultured in 6-well format in FGM. At ∼80% confluency, cells were treated with the indicated drugs or switched into amino acid-free media (USBiological, #D9800-27) for 6h. Cells were washed with ice cold HBS twice and plates were snap frozen in liquid nitrogen. Metabolite extraction was performed as described previously^83^. In brief, wells with the same number of cells were extracted using ice-cold 80% methanol in HPLC-grade water with 4 μg/mL norvaline internal standard (Sigma-Aldrich N7627). After centrifuging the extracts at 21,000xG for 10m at 4°C, the soluble supernatant was transferred to a fresh tube and dried under nitrogen gas. Polar metabolites were derivatized by resuspending the dried sample in 24μL of 2% methoxylamine hydrochloride in pyridine (MOX; ThermoFisher #TS–45950) at 37°C for 1h. 30μL N–methyl–N–(tert– butyldimethylsilyl) trifluoroacetamide +1% tert–Butyldimethylchlorosilane (Sigma-Aldrich #375934) was added and samples were heated to 80°C for 2h. The derivatized samples were analyzed on a DB-35MS column (30 m × 0.25 mm i.d. × 0.25 μm, Agilent J&W Scientific) on an Agilent 7890B gas chromatograph/Agilent 5977B mass spectrometer. Metabolite ion counts were quantified by integrating GC-MS ion fragments for each amino acid derivative (El Maven software v11.0, Elucidata), corrected for natural isotope abundance using the R package IsoCorrectoR, and normalized to the internal norvaline standard.

### *In silico* molecular screening and docking

*In silico* molecular screening and docking of the inhibitors within GCN2 and BRAF were completed with the Schrödinger 2024-2 Suite, using the Glide module^84^. The RAF inhibitors were prepared along with BRAF^V600E^ and GCN2 wild-type crystal structures (PDB code: [4MNF] and [6N3O], respectively). The FDA-approved & Passed Phase I Drug Library from Selleckchem was also prepared. The library was screened against the prepared GCN2 structure for relative ranking along with MM/GBSA ligand-protein energies. The RAF inhibitors were docked into the BRAF and GCN2 structures at ATP active site (along with ATP as a control) with Standard and Extra Precision (sorafenib was docked into CRAFWT, PDB code: [3OMV], instead of BRAF). Two-dimensional plots, surfaces, and corresponding MM/GBSA energies were then generated.

## References

1. Miller, K. D. et al. Cancer treatment and survivorship statistics, 2022. CA: A Cancer Journal for Clinicians 72, 409–436 (2022).

2. Hata, A. N. et al. Tumor cells can follow distinct evolutionary paths to become resistant to epidermal growth factor receptor inhibition. Nat Med 22, 262–269 (2016).

3. Sharma, S. V. et al. A Chromatin-Mediated Reversible Drug-Tolerant State in Cancer Cell Subpopulations. Cell 141, 69–80 (2010).

4. Arozarena, I. & Wellbrock, C. Phenotype plasticity as enabler of melanoma progression and therapy resistance. Nat Rev Cancer 19, 377–391 (2019).

5. Boumahdi, S. & de Sauvage, F. J. The great escape: tumour cell plasticity in resistance to targeted therapy. Nat Rev Drug Discov 19, 39–56 (2020).

6. Su, Y. et al. Single-cell analysis resolves the cell state transition and signaling dynamics associated with melanoma drug-induced resistance. Proc Natl Acad Sci U S A 114, 13679–13684 (2017).

7. Siegel, R. L., Giaquinto, A. N. & Jemal, A. Cancer statistics, 2024. CA Cancer J Clin 74, 12–49 (2024).

8. Wagle, N. et al. Dissecting therapeutic resistance to RAF inhibition in melanoma by tumor genomic profiling. J Clin Oncol 29, 3085–3096 (2011).

9. Long, G. V. et al. Increased MAPK reactivation in early resistance to dabrafenib/trametinib combination therapy of BRAF-mutant metastatic melanoma. Nat Commun 5, 5694 (2014).

10. Davies, H. et al. Mutations of the BRAF gene in human cancer. Nature 417, 949–954 (2002).

11. Kim, A. & Cohen, M. S. The discovery of vemurafenib for the treatment of BRAF-mutated metastatic melanoma. Expert Opin Drug Discov 11, 907–916 (2016).

12. Rheault, T. R. et al. Discovery of Dabrafenib: A Selective Inhibitor of Raf Kinases with Antitumor Activity against B-Raf-Driven Tumors. ACS Med Chem Lett 4, 358–362 (2013).

13. Okten, I. N., Ismail, S., Withycombe, B. M. & Eroglu, Z. Preclinical discovery and clinical development of encorafenib for the treatment of melanoma. Expert Opin Drug Discov 15, 1373–1380 (2020).

14. Grippo, J. F. et al. A phase I, randomized, open-label study of the multiple-dose pharmacokinetics of vemurafenib in patients with BRAFV600Emutation-positive metastatic melanoma. Cancer Chemother Pharmacol 73, 103–111 (2014).

15. Waddell, J. A. & Solimando, D. A. Drug Monographs: Dabrafenib and Trametinib. Hosp Pharm 48, 818–821 (2013).

16. Sullivan, R. J. et al. A Phase Ib/II Study of the BRAF Inhibitor Encorafenib Plus the MEK Inhibitor Binimetinib in Patients with BRAFV600E/K-mutant Solid Tumors. Clinical Cancer Research 26, 5102–5112 (2020).

17. Yang, C., Tian, C., Hoffman, T. E., Jacobsen, N. K. & Spencer, S. L. Melanoma subpopulations that rapidly escape MAPK pathway inhibition incur DNA damage and rely on stress signalling. Nat Commun 12, 1747 (2021).

18. Goyal, Y. et al. Diverse clonal fates emerge upon drug treatment of homogeneous cancer cells. Nature 620, 651–659 (2023).

19. Hoffman, T. E. et al. Multiple cancers escape from multiple MAPK pathway inhibitors and use DNA replication stress signaling to tolerate aberrant cell cycles. Science Signaling 16, eade8744 (2023).

20. Lines, C. L., McGrath, M. J., Dorwart, T. & Conn, C. S. The integrated stress response in cancer progression: a force for plasticity and resistance. Front. Oncol. 13, (2023).

21. Dong, J., Qiu, H., Garcia-Barrio, M., Anderson, J. & Hinnebusch, A. G. Uncharged tRNA activates GCN2 by displacing the protein kinase moiety from a bipartite tRNA-binding domain. Mol Cell 6, 269–279 (2000).

22. Han, A. P. et al. Heme-regulated eIF2alpha kinase (HRI) is required for translational regulation and survival of erythroid precursors in iron deficiency. EMBO J 20, 6909–6918 (2001).

23. Fessler, E. et al. A pathway coordinated by DELE1 relays mitochondrial stress to the cytosol. Nature 579, 433–437 (2020).

24. Guo, X. et al. Mitochondrial stress is relayed to the cytosol by an OMA1-DELE1-HRI pathway. Nature 579, 427–432 (2020).

25. Kalkavan, H. et al. Sublethal cytochrome c release generates drug-tolerant persister cells. Cell 185, 3356–3374.e22 (2022).

26. Sekine, Y. et al. A mitochondrial iron-responsive pathway regulated by DELE1. Mol Cell 83, 2059–2076.e6 (2023).

27. Balachandran, S. et al. Essential role for the dsRNA-dependent protein kinase PKR in innate immunity to viral infection. Immunity 13, 129–141 (2000).

28. Shi, Y. et al. Identification and characterization of pancreatic eukaryotic initiation factor 2 alpha-subunit kinase, PEK, involved in translational control. Mol Cell Biol 18, 7499–7509 (1998).

29. Harding, H. P., Zhang, Y. & Ron, D. Protein translation and folding are coupled by an endoplasmic-reticulum-resident kinase. Nature 397, 271–274 (1999).

30. Yang, W. & Hinnebusch, A. G. Identification of a regulatory subcomplex in the guanine nucleotide exchange factor eIF2B that mediates inhibition by phosphorylated eIF2. Mol Cell Biol 16, 6603–6616 (1996).

31. Krishnamoorthy, T., Pavitt, G. D., Zhang, F., Dever, T. E. & Hinnebusch, A. G. Tight binding of the phosphorylated alpha subunit of initiation factor 2 (eIF2alpha) to the regulatory subunits of guanine nucleotide exchange factor eIF2B is required for inhibition of translation initiation. Mol Cell Biol 21, 5018–5030 (2001).

32. Vattem, K. M. & Wek, R. C. Reinitiation involving upstream ORFs regulates ATF4 mRNA translation in mammalian cells. Proceedings of the National Academy of Sciences 101, 11269–11274 (2004).

33. Pakos-Zebrucka, K. et al. The integrated stress response. EMBO Rep 17, 1374–1395 (2016).

34. Neill, G. & Masson, G. R. A stay of execution: ATF4 regulation and potential outcomes for the integrated stress response. Frontiers in Molecular Neuroscience 16, (2023).

35. Wortel, I. M. N., van der Meer, L. T., Kilberg, M. S. & van Leeuwen, F. N. Surviving Stress: Modulation of ATF4-Mediated Stress Responses in Normal and Malignant Cells. Trends Endocrinol Metab 28, 794–806 (2017).

36. Moeckel, S. et al. ATF4 contributes to autophagy and survival in sunitinib treated brain tumor initiating cells (BTICs). Oncotarget 10, 368–382 (2019).

37. Pike, L. R. G. et al. Transcriptional up-regulation of ULK1 by ATF4 contributes to cancer cell survival. Biochem J 449, 389–400 (2013).

38. B’chir, W. et al. The eIF2α/ATF4 pathway is essential for stress-induced autophagy gene expression. Nucleic Acids Res 41, 7683–7699 (2013).

39. Matsumoto, H. et al. Selection of autophagy or apoptosis in cells exposed to ER-stress depends on ATF4 expression pattern with or without CHOP expression. Biol Open 2, 1084–1090 (2013).

40. Nagasawa, I., Kunimasa, K., Tsukahara, S. & Tomida, A. BRAF-mutated cells activate GCN2-mediated integrated stress response as a cytoprotective mechanism in response to vemurafenib. Biochem Biophys Res Commun 482, 1491–1497 (2017).

41. Armstrong, J. L., Flockhart, R., Veal, G. J., Lovat, P. E. & Redfern, C. P. F. Regulation of endoplasmic reticulum stress-induced cell death by ATF4 in neuroectodermal tumor cells. J Biol Chem 285, 6091–6100 (2010).

42. Bou-Nader, C. et al. Gcn2 structurally mimics and functionally repurposes the HisRS enzyme for the integrated stress response. Proceedings of the National Academy of Sciences 121, e2409628121 (2024).

43. Gold, L. T. & Masson, G. R. GCN2: roles in tumour development and progression. Biochem Soc Trans 50, 737–745 (2022).

44. Inglis, A. J. et al. Activation of GCN2 by the ribosomal P-stalk. Proceedings of the National Academy of Sciences 116, 4946–4954 (2019).

45. Darnell, A. M., Subramaniam, A. R. & O’Shea, E. K. Translational Control through Differential Ribosome Pausing during Amino Acid Limitation in Mammalian Cells. Mol Cell 71, 229–243.e11 (2018).

46. Ye, J. et al. GCN2 sustains mTORC1 suppression upon amino acid deprivation by inducing Sestrin2. Genes Dev 29, 2331–2336 (2015).

47. Ge, M.-K. et al. The tRNA-GCN2-FBXO22-axis-mediated mTOR ubiquitination senses amino acid insufficiency. Cell Metabolism 35, 2216–2230.e8 (2023).

48. Darawshi, O. et al. Phosphorylation of GCN2 by mTOR confers adaptation to conditions of hyper-mTOR activation under stress. Journal of Biological Chemistry 300, (2024).

49. Torrence, M. E. et al. The mTORC1-mediated activation of ATF4 promotes protein and glutathione synthesis downstream of growth signals. eLife 10, e63326 (2021).

50. Jiang, H.-Y. & Wek, R. C. GCN2 phosphorylation of eIF2α activates NF-κB in response to UV irradiation. Biochem J 385, 371–380 (2005).

51. Deng, J. et al. Activation of GCN2 in UV-Irradiated Cells Inhibits Translation. Current Biology 12, 1279–1286 (2002).

52. Alasiri, G. et al. Reciprocal regulation between GCN2 (eIF2AK4) and PERK (eIF2AK3) through the JNK-FOXO3 axis to modulate cancer drug resistance and clonal survival. Mol Cell Endocrinol 515, 110932 (2020).

53. Nakamura, A. et al. Inhibition of GCN2 sensitizes ASNS-low cancer cells to asparaginase by disrupting the amino acid response. Proceedings of the National Academy of Sciences 115, E7776–E7785 (2018).

54. Ghosh, J. C. et al. Ghost mitochondria drive metastasis through adaptive GCN2/Akt therapeutic vulnerability. Proceedings of the National Academy of Sciences 119, e2115624119 (2022).

55. Drees, J. et al. 1327 Novel GCN2 modulator HC-7366 decreases pulmonary metastases and reduces myeloid-derived suppressor cells. J Immunother Cancer 10, (2022).

56. Örd, T., Örd, D., Adler, P. & Örd, T. Genome-wide census of ATF4 binding sites and functional profiling of trait-associated genetic variants overlapping ATF4 binding motifs. PLoS Genet 19, e1011014 (2023).

57. Tang, C. P. et al. GCN2 kinase activation by ATP-competitive kinase inhibitors. Nat Chem Biol 18, 207–215 (2022).

58. Szaruga, M. et al. Activation of the integrated stress response by inhibitors of its kinases. Nat Commun 14, 5535 (2023).

59. Carlson, K. R., Georgiadis, M. M., Tameire, F., Staschke, K. A. & Wek, R. C. Activation of Gcn2 by small molecules designed to be inhibitors. J Biol Chem 299, 104595 (2023).

60. Drainas, A. P. et al. GCN2 is a determinant of the response to WEE1 kinase inhibition in small-cell lung cancer. Cell Rep 43, 114606 (2024).

61. Liang, S.-H., Zhang, W., Mcgrath, B. C., Zhang, P. & Cavener, D. R. PERK (eIF2α kinase) is required to activate the stress-activated MAPKs and induce the expression of immediate-early genes upon disruption of ER calcium homoeostasis. Biochem J 393, 201–209 (2006).

62. Spencer, S. L. et al. The Proliferation-Quiescence Decision Is Controlled by a Bifurcation in CDK2 Activity at Mitotic Exit. Cell 155, 369–383 (2013).

63. Tian, C., Yang, C. & Spencer, S. L. EllipTrack: A Global-Local Cell-Tracking Pipeline for 2D Fluorescence Time-Lapse Microscopy. Cell Rep 32, 107984 (2020).

64. Wong, Y. L. et al. eIF2B activator prevents neurological defects caused by a chronic integrated stress response. eLife 8, e42940.

65. Puthenveetil, S. et al. Controlling activation of the RNA-dependent protein kinase by siRNAs using site-specific chemical modification. Nucleic Acids Res 34, 4900–4911 (2006).

66. Cottrell, K. A. et al. Activation of PKR by a short-hairpin RNA. Sci Rep 14, 23533 (2024).

67. Ye, J. et al. The GCN2-ATF4 pathway is critical for tumour cell survival and proliferation in response to nutrient deprivation. The EMBO Journal 29, 2082–2096 (2010).

68. Eckert, S. et al. Decrypting the molecular basis of cellular drug phenotypes by dose-resolved expression proteomics. Nat Biotechnol 1–10 (2024) doi:10.1038/s41587-024-02218-y.

69. Llargués-Sistac, G., Bonjoch, L. & Castellvi-Bel, S. HAP1, a new revolutionary cell model for gene editing using CRISPR-Cas9. Front Cell Dev Biol 11, 1111488 (2023).

70. Tauber, D. & Parker, R. 15-Deoxy-Δ12,14-prostaglandin J2 promotes phosphorylation of eukaryotic initiation factor 2α and activates the integrated stress response. J Biol Chem 294, 6344–6352 (2019).

71. Genheden, S. & Ryde, U. The MM/PBSA and MM/GBSA methods to estimate ligand-binding affinities. Expert Opin Drug Discov 10, 449–461 (2015).

72. Togashi, Y. et al. Pharmacokinetics of Erlotinib and Its Active Metabolite OSI-420 in Patients with Non-small Cell Lung Cancer and Chronic Renal Failure Who Are Undergoing Hemodialysis. Journal of Thoracic Oncology 5, 601–605 (2010).

73. Keyvanjah, K. et al. Pharmacokinetics of neratinib during coadministration with lansoprazole in healthy subjects. Br J Clin Pharmacol 83, 554–561 (2017).

74. Lin, A. et al. Off-target toxicity is a common mechanism of action of cancer drugs undergoing clinical trials. Sci Transl Med 11, eaaw8412 (2019).

75. Shyam Sunder, S., Sharma, U. C. & Pokharel, S. Adverse effects of tyrosine kinase inhibitors in cancer therapy: pathophysiology, mechanisms and clinical management. Sig Transduct Target Ther 8, 1–27 (2023).

76. Gilley, R. et al. RAF inhibitors activate the integrated stress response by direct activation of GCN2. 2024.08.15.607884 Preprint at 10.1101/2024.08.15.607884 (2024).

77. Neill, G. et al. Paradoxical Activation of GCN2 by ATP-competitive inhibitors via allosteric activation and autophosphorylation. 2024.08.14.606984 Preprint at 10.1101/2024.08.14.606984 (2024).

78. Castilho, B. A. et al. Keeping the eIF2 alpha kinase Gcn2 in check. Biochim Biophys Acta 1843, 1948–1968 (2014).

79. Grob, J. J. et al. Comparison of dabrafenib and trametinib combination therapy with vemurafenib monotherapy on health-related quality of life in patients with unresectable or metastatic cutaneous BRAF Val600-mutation-positive melanoma (COMBI-v): results of a phase 3, open-label, randomised trial. Lancet Oncol 16, 1389–1398 (2015).

80. Rasco, D. W. et al. Phase 1 study of the pan-RAF inhibitor tovorafenib in patients with advanced solid tumors followed by dose expansion in patients with metastatic melanoma. Cancer Chemother Pharmacol 92, 15–28 (2023).

81. Liu, Y., Azizian, N. G., Sullivan, D. K. & Li, Y. mTOR inhibition attenuates chemosensitivity through the induction of chemotherapy resistant persisters. Nat Commun 13, 7047 (2022).

82. Panwar, V. et al. Multifaceted role of mTOR (mammalian target of rapamycin) signaling pathway in human health and disease. Sig Transduct Target Ther 8, 1–25 (2023).

83. Rong, Y., Darnell, A. M., Sapp, K. M., Vander Heiden, M. G. & Spencer, S. L. Cells use multiple mechanisms for cell-cycle arrest upon withdrawal of individual amino acids. Cell Rep 42, 113539 (2023).

84. Friesner, R. A. et al. Glide: A New Approach for Rapid, Accurate Docking and Scoring. 1. Method and Assessment of Docking Accuracy. J. Med. Chem. 47, 1739–1749 (2004).

