## Supplemental Figures for "BRAF^V600^ and ErbB inhibitors directly activate GCN2 in an off-target manner to limit cancer cell proliferation"

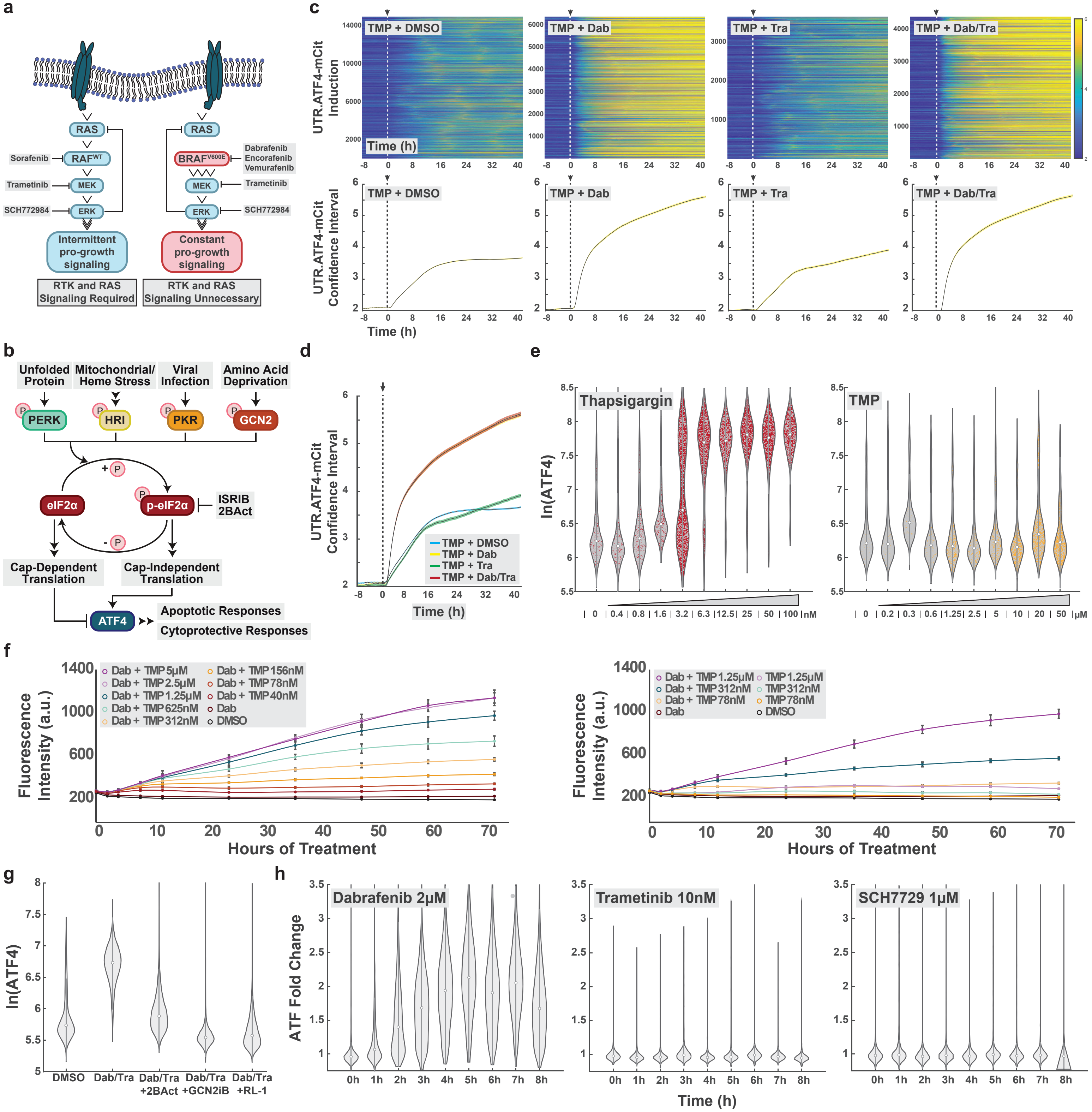


**Figure S1. a** Schematic of canonical (left) and BRAF-mutant (right) MAPK signaling. The drugs utilized herein are indicated next to their canonical target. **b** Schematic of canonical ISR signaling. **c** Activation of the UTR.ATF4-mCit sensor in response to the indicated rugs (TMP 1μM; dabrafenib 2μM; trametinib 10nM). Single-cell traces over time are displayed as heatmaps (top, reproduced from Fig. 1b). Confidence interval plots (bottom) show mean sensor activation (black) and 95% confidence interval (yellow) over time. **d** Overlay of 95% confidence interval plots from (c). The red and yellow curves lie precisely on top of one another and appear as orange. **e** A375 cells were treated with increasing doses of thapsigargin (tg, a canonical PERK activator) or TMP for 6h. Single-cell immunofluorescence of ATF4 is plotted as violins, revealing no activation of the ISR by TMP. **f** A375 cells containing the UTR.ATF4-mCit sensor and H2B-mCherry were subjected to TMP titration with dabrafenib (left, 2μM). The effects of TMP during activating conditions (plus dabrafenib) vs. non-activating conditions (TMP alone) are compared on the right. Mean sensor activity across 4 wells was plotted. Error bars represent +/- 1 standard deviation. **g** A375 cells were treated with 2μM dabrafenib plus 10nM trametinib and the indicated drug for 6h, preceded by 1h pretreatment with 2BAct (200nM), GCN2iB (1μM) or Rapalink-1 (1μM). Single-cell immunofluorescence of ATF4 is plotted as violins. **h** A375 cells were treated with the indicated drug over 8h. Single-cell immunofluorescence of ATF4 is plotted as violins, showing fold change over 0h (DMSO).

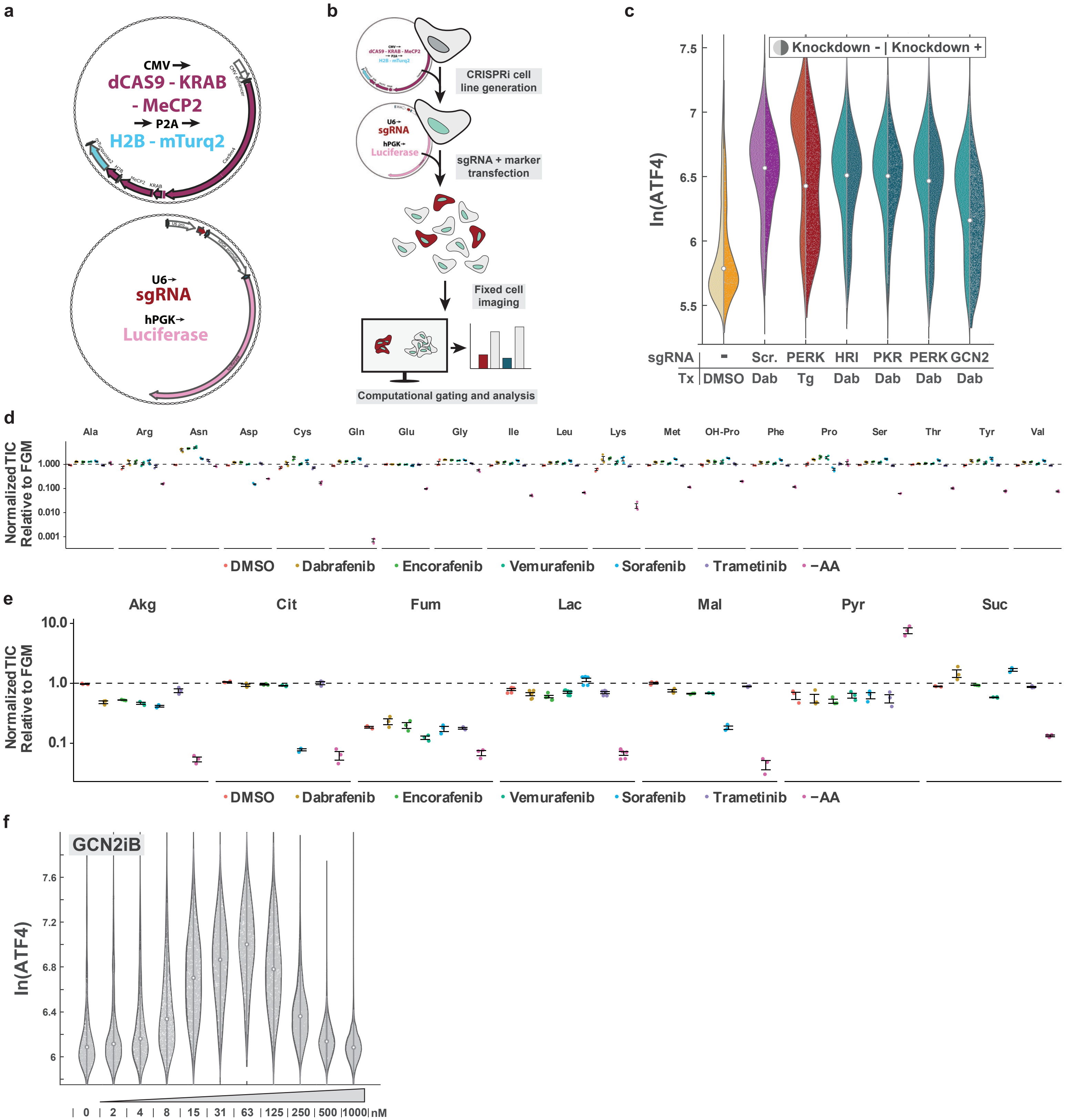


**Figure S2. a** Representations of the plasmids used in the scCRISPRi system. **b** Schematic of the developed single cell CRISPRi (scCRISPRi) knockdown platform (see Methods). **c** CRISPRi knockdown data performed in A375 cells with the indicated ISR kinase knockdown target or scramble control (Scr.). Dabrafenib was utilized at 2μM, and thapsigargin (tg) at 1μM. Data are from the same experiment as Fig. 2a, but here we directly compare ATF4 levels in cells that did not receive the knockdown (left side of split violin) vs. cells that did receive the knockdown (right side of split violin) from the same wells of the 96-well plate. Single-cell immunofluorescence of mean nuclear ATF4 signal is plotted. **d** A375 cells were treated with the indicated drug or treatment for 6h in biological triplicate (dabrafenib 2μM, encorafenib 2μM, vemurafenib 32μM, sorafenib 16μM, trametinib 10nM). Gas chromatography - mass spectrometry was performed and amino acid abundance was quantified. Error bars represent +/- 1 standard deviation. **e** Same as in (d), but central carbon metabolite abundance was quantified. **f** Single-cell immunofluorescence of ATF4 in A375 cells is plotted as violins for increasing doses of GCN2iB treatment for 6h.

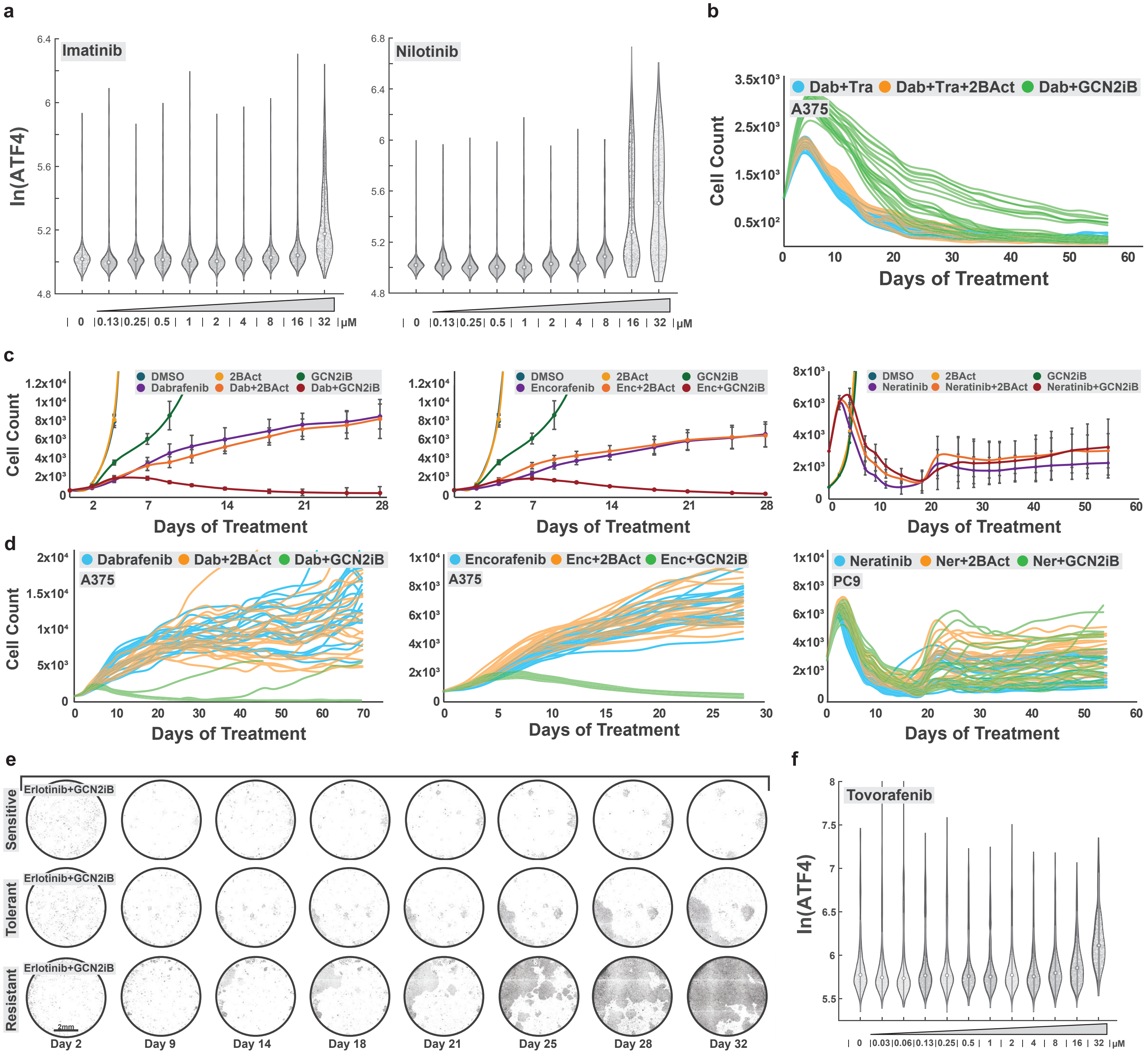

**Figure S3. a** A375 cells were subjected to increasing concentrations of imatinib (left) or nilotinib (right) for 6h. Single-cell immunofluorescence of mean nuclear ATF4 signal is plotted. **b** Single well traces of A375 cells treated with dabrafenib (2μM) + trametinib (10nM) and GCN2iB (1 μM) or 2BAct (200nM) corresponding to the outgrowth in Fig. 5b (right). **c** The same cell count experiment as in Fig. 5b were performed in A375 cells with dabrafenib (2μM) (cotreatments GCN2iB 1μM; 2BAct 200nM) or encorafenib (2μM), or in PC9 cells with neratinib (500nM, Cmax 152nM) (cotreatments GCN2iB 1μM; 2BAct 200nM). Cell count was quantified. Error bars represent +/- 1 standard deviation. **d** Single well traces of the cell counts shown in (c). **e** Representative full well images of PC9 H2B-mCherry cells in the sensitive, tolerant, and resistant states up to day 32 of erlotinib (5μM) + GCN2iB (1μM) cotreatment. **f** A375 cells were treated with increasing doses of tovorafenib for 6h. Single-cell immunofluorescence of mean nuclear ATF4 signal is plotted as violins.
